# SABRE populates ER domains essential for cell plate maturation and cell expansion influencing cell and tissue patterning

**DOI:** 10.1101/2020.11.24.396788

**Authors:** Xiaohang Cheng, Magdalena Bezanilla

## Abstract

The SABRE protein, originally identified in plants, is found throughout eukaryotes. In plants, SABRE has been implicated in cell expansion, division plane orientation and planar polarity. However, how SABRE mediates these processes remains an open question. Here, we have taken advantage of the fact that the bryophyte *Physcomitrium patens* has a single copy of *SABRE*, is an excellent model for cell biology and is readily amenable to precise genetic alterations to investigate SABRE’s mechanism of action. We discovered that SABRE null mutants were stunted in both polarized growing and diffusely growing tissues, similar to reported phenotypes in seed plants. However, in polarized growing cells, we observed significant delays in cell plate formation and sometimes catastrophic failures in cell division. We generated a functional SABRE fluorescent fusion protein and determined that it forms dynamic puncta on regions of the endoplasmic reticulum (ER) both in the cytoplasm during interphase and at the new cell plate during division. In the absence of *SABRE*, ER morphology was severely compromised with large aggregates accumulating in the cytoplasm and abnormal buckling along the developing cell plate late in cytokinesis. In fact, SABRE and the ER maximally accumulated on the developing plate specifically during cell plate maturation, coincident with the timing of the onset of failures in cell plate formation in cells lacking SABRE. Further we discovered that callose deposition is delayed in *Δsabre* cells, and in cells that failed to divide, abnormal callose accumulations formed at the cell plate. Our findings demonstrated that SABRE functions by influencing the ER and callose deposition, revealing a surprising and essential role for the ER in cell plate maturation. Given that SABRE is conserved, understanding how SABRE influences cell and tissue patterning has profound significance across eukaryotes.

## Introduction

Due to their sessile nature, plants can’t simply run away from environmental stimuli and instead must adjust to their environment by regulating growth patterns. Plant growth is a coupled process involving deposition of extra cellular matrix material – the cell wall – around individual cells and cell expansion. Precise regulation of the composition of this matrix ensures where a particular cell can expand. To control cell shape throughout a tissue, polarity cues at the cellular and tissue level ensures coordination ultimately patterning whole organs, such as correctly oriented roots and stems (Blilou et al., 2005; Kania et al., 2014; van Dop et al., 2020), as well as specialized structures including stomata and root hairs (Gilroy and Jones, 2000; Houbaert et al., 2018; Mansfield et al., 2018; Zhang et al., 2016). Positioning the cell division plane contributes to cell shape and provides polarity information (Zhang and Dong, 2018). For example, in root and shoot tissue in seed plants, the cell division plane is perpendicular to the growth axis and creates polygonal cells that are aligned longitudinally with each other (Ambrose et al., 2007; Galjart, 2005; Schaefer et al., 2017; Smith et al., n.d.; Walker et al., 2007), ensuring that expansion is aligned uniformly with the overall plant growth axis. For specialized cell types, polarized cell wall deposition and cell plate positioning cooperates to determine cell morphology. In the leaf epidermis, asymmetric cell divisions define the stomatal guard cells (Dong et al., 2009), and restricted expansion in cells define the jigsaw-shaped epidermal cells (Sapala et al., 2019). In filamentous cells such as pollen tubes and root hairs in seed plants, polarized secretion of flexible wall material to the cell apex leads to cell expansion occurring only at the apex of the cell (Bascom et al., 2018; Chen et al., 2018; Dehors et al., 2019; Orr et al., 2020).

Using forward genetics, many studies have identified mutations that alter plant morphogenesis and have provided insights into the regulation of expansion and polarity, demonstrating that plants regulate cell wall deposition in a myriad of ways. However, in some cases it has been challenging to molecularly link how the proteins identified by these mutant screens effect their influence on morphogenesis. The *sabre* mutant, which has short fat roots, was first identified in Arabidopsis in the early 1990s (Benfey et al., 1993). The increased root diameter resulted from exaggerated radial expansion primarily in root cortex cells, suggesting that SABRE plays a role in regulating expansion of diffusely growing cells (Aeschbacher et al., 1995). A second copy of *SABRE*, named *KINKY POLLEN (KIP)*, which is expressed most strongly in roots, pollen and developing seeds, was identified in a screen for abnormal pollen tube and root hair morphology (Procissi et al., 2003). Plants lacking KIP form defective pollen tubes that are characterized by periods of relatively normal growth punctuated with periods of slow growth and growth arrests. Recovery from these growth arrests often led to growth initiating in new directions, ultimately leading to the kinky or twisty phenotype. In addition, root hairs were shorter and thicker in *kip* mutants (Procissi et al., 2003). These data suggest that SABRE contributes to diffuse growth while KIP contributes to polarized growth. However, the homozygous *kip/sab* double mutant exhibited enhanced phenotypes in both diffuse and polarized growing tissues, indicating overlapping function of these two homologs (Procissi et al., 2003). More recent studies have found that in *sabre* mutants, cell plate positioning in the root meristem was variable, resulting in cells that were not cylindrically aligned. Furthermore, root hair emergence was no longer restricted to the basal portion of the epithelial cell (Pietra et al., 2013). In *sabre*, transcription factors that initiate root hair cell fate were also altered, resulting in the formation of root hairs from ectopic sites. Collectively these studies point to a critical role for *SABRE* in regulating plant polarity at both the cell and tissue levels (Pietra et al., 2015).

In *Zea mays, ABERRANT POLLEN TRANSMISSION 1 (APT1)* gene was identified as the *SABRE/KIP* homolog, whose mutation also resulted in short and meandering pollen tubes (Xu and Dooner, 2006). Consistent with predictions of a Golgi localization sequence at the C-terminus of SABRE homologs (Pietra et al., 2013; Xu and Dooner, 2006), expressing C-terminal fragments of APT1 fused with fluorescent proteins in tobacco pollen tubes resulted in localization to the Golgi. However, full length SABRE stably expressed in Arabidopsis exhibited punctate localization in the cytosol of root epidermal cells that did not obviously represent any known endomembrane compartment (Pietra et al., 2013). More detailed localization studies are needed to help reconcile these apparently contradictory findings.

Despite studies characterizing *SABRE* gene function at the physiological level, and protein localization at the cellular level, the molecular basis of SABRE function remains unclear. Interestingly, the moss *Physcomitrium* (*Physcomitrella*) *patens* has a single copy of *SABRE. P. patens* is amenable to rapid CRISPR-Cas9 mediated genome editing (Collonnier et al., 2017; Lopez-Obando et al., 2016; Mallett et al., 2019). Plants can be propagated asexually indefinitely, avoiding potential problems resulting from defects in sexual reproduction. Additionally, *P. patens* is an ideal system to study how cell shape affects developmental patterning (Rensing et al., 2020; Rounds and Bezanilla, 2013). Moss juvenile tissue, protonemata, is haploid and comprised of a filamentous two-dimensional branching network that is a single-cell layer thick, making it readily amenable to high resolution microscopy. Protonemal tissue expands exclusively by polarized growth with cell division occurring in the apical cell of the filament. Subapical cells re-enter the cell cycle once a protrusion emerges generating a new filament, with the branching cell dividing at the base of the emerging protrusion. As protonemata age, some of the protrusions switch fates to bud-like structures that expand in three dimensions, ultimately resulting in the development of leafy shoots, known as gametophores, the adult tissues.

Here, using genome editing we generated a null *sabre* mutant and functional fluorescent fusions at the native genomic locus to investigate the molecular basis of *SABRE* function in *P. patens*. We found that *Δsabre* plants are stunted, exhibiting defects in polarized growth, diffuse cell expansion and dramatic cell division failures accompanied by deposition of brown material into the cytoplasm often resulting in cell death. Surprisingly, even with the polarized growth and division defects, SABRE did not localize to the cytoskeleton. Instead, SABRE localized to a fraction of the ER near the tip of the cell, at the cell cortex, and in the phragmoplast during cell division. These results indicate that SABRE regulates plant cell expansion and division via its interaction with the ER, pointing to a fundamentally important role for the ER in plant cell and tissue morphogenesis.

## Results

### Loss of SABRE function inhibits polarized growth and diffuse cell expansion

The *P. patens* genome has one *SABRE* gene that encodes for a 2736 aa protein. To disrupt *SABRE*, we used CRISPR-Cas9 mediated homology directed repair to insert a 363 bp cassette with stop codons in all three possible frames into exon 2 of the *SABRE* locus (Figure 1 – figure supplement 1). The mutant allele results in a frame shift starting at amino acid 38 followed by a premature stop codon after amino acid 47. For consistency, all *Δsabre* lines generated in this study contained the exact same mutation. Compared to wild type, *Δsabre* plants had smaller and more compact protonemata (Figure 1A). To quantify this difference, we regenerated plants from single protoplasts and measured the overall size seven days after protoplasting (Figure 1B). Compared to protonemata in control plants, we found that *Δsabre* protonemata were 60% smaller (Figure 1C).

**Figure 1.**
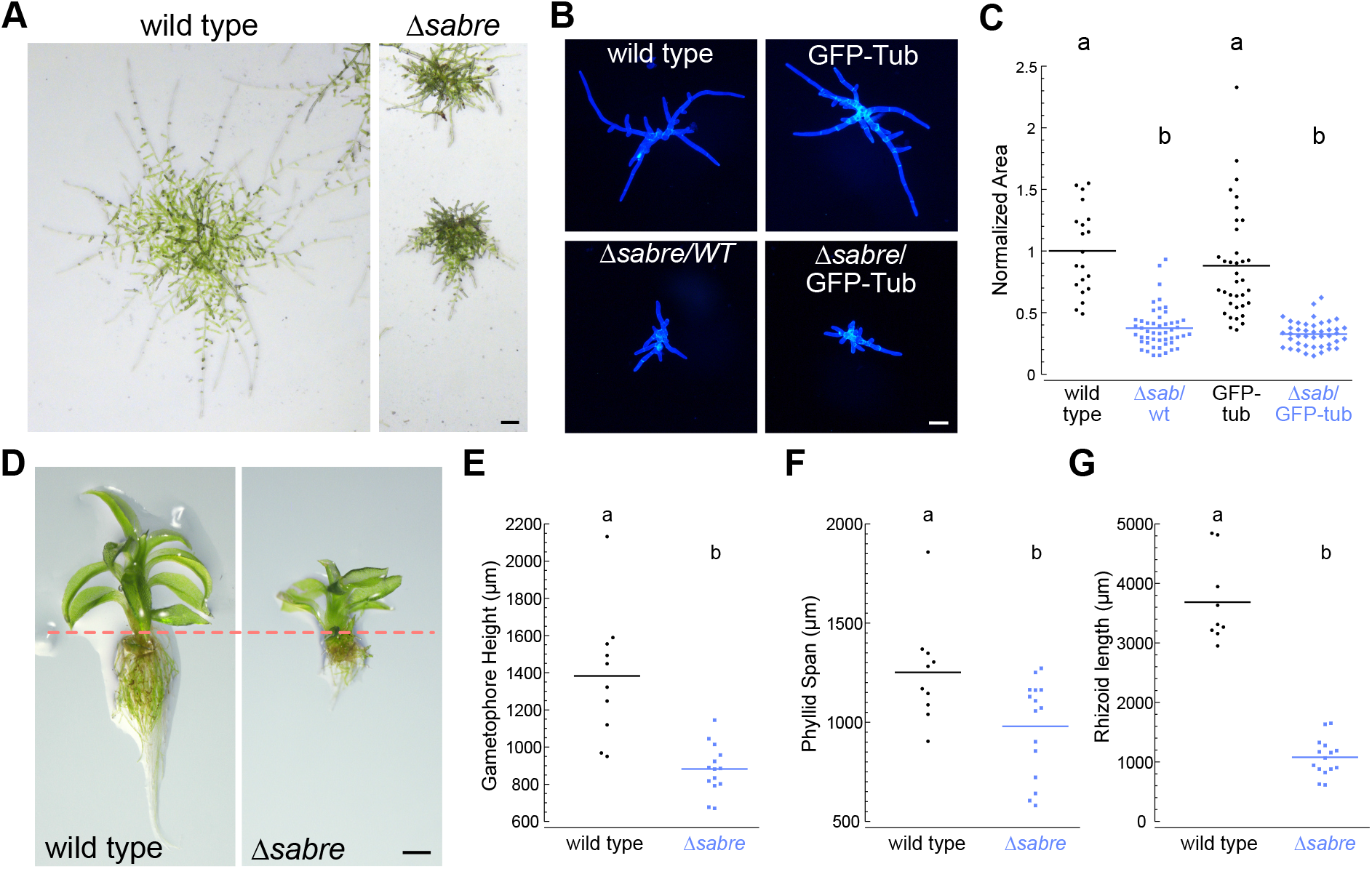
*SABRE* influences both polarized cell expansion and diffuse growth. (A) Example images of two week-old wild type and *Δsabre* plants regenerated from single protoplasts. Extended-depth-of-focus (EDF) images were created from Z-stacks acquired with a stereo microscope. Scale bar, 200 μm. (B) Representative fluorescence images of 7 day-old plants regenerated from protoplasts. Images of plants stained with calcofluor white were acquired with a fluorescent stereo microscope. Scale bar, 100 μm. (C) Quantification of *Δsabre* plant size calculated from area of calcofluor fluorescence. Plant area was normalized to wild type. N=20, wild type; N=50, Δsabre/WT; N=36, GFP-tub; N=43 Δsabre/GFP-tub. Letters indicate groups with significantly different means as determined by ANOVA with a Tukey’s HSD all pair comparison post-hoc test (α=0.05). (D) Extended depth-of-focus (EDF) images of example mature gametophores. Scale bar, 500 μm. Dashed line indicates the boundary between the aerial tissue (top) and the rhizoids (bottom). (E-G) Quantification of gametophore height, phyllid span and rhizoid length. Letters indicate groups with significantly different means determined by Student’s t-test for unpaired data with equal variance. (E) N=10, wild type; N=14, *Δsabre*. (F) N=10, wild type; N=15, *Δsabre*. (G) N=9, wild type; N=15, *Δsabre*.

While protonemal filaments increase in size exclusively by polarized expansion at the tip of the apical cell, older tissue produces gametophores that expand by diffuse growth (Figure 1D). We also observed that *Δsabre* gametophores are 36% shorter than wild type (Figure 1D, E). The phyllids, leaf-like structures that emanate from the gametophore, were 22% smaller than wild type (Figure 1F). Rhizoids are polarized-growing filaments that grow from the base of the gametophore anchoring it in the soil. *Δsabre* rhizoids were 70% shorter than wild type rhizoids, a decrease in size comparable to the polarized-growing protonemata (Figure 1G). Time-lapse imaging demonstrated that while early developmental patterning of *Δsabre* gametophores was not altered, cell expansion was significantly delayed (Figure 1 – figure supplement 2, video 1). Together, these data demonstrate that the single *SABRE* gene regulates both polarized and diffuse growth in *P. patens*.

To determine whether smaller organ size resulted from changes in underlying cell size, we imaged protonemata and gametophore development at higher resolution (Figure 2). During protonemal development, the apical stem cell expands by polarized growth and divides, leaving behind a subapical cell that does not elongate anymore. Thus, we measured the length of protonemal subapical cells. While *Δsabre* cells were 30% shorter than wild type (Figure 2A), the decrease of overall plant size was 60% (Figure 1C). This discrepancy could be due to defects in the rate of growth or how often a particular filament is actively growing. To distinguish between these two possibilities, we measured cell expansion rates in actively growing apical cells as determined by time-lapse imaging. While on average actively growing *Δsabre* cells grew only 20% slower than control cells, the reduction in growth was not significantly different. In contrast, time-lapse imaging revealed that over the same time period *Δsabre* protonemal filaments often grew significantly less than a comparable wild type filament due to the fact that for a large portion of the time-lapse acquisition the *Δsabre* cell was not growing (Figure 2B – video 2). As a result, we reasoned that additional mechanisms, such as frequent long pauses in growth, likely contributed to the decrease in plant size.

**Figure 2.**
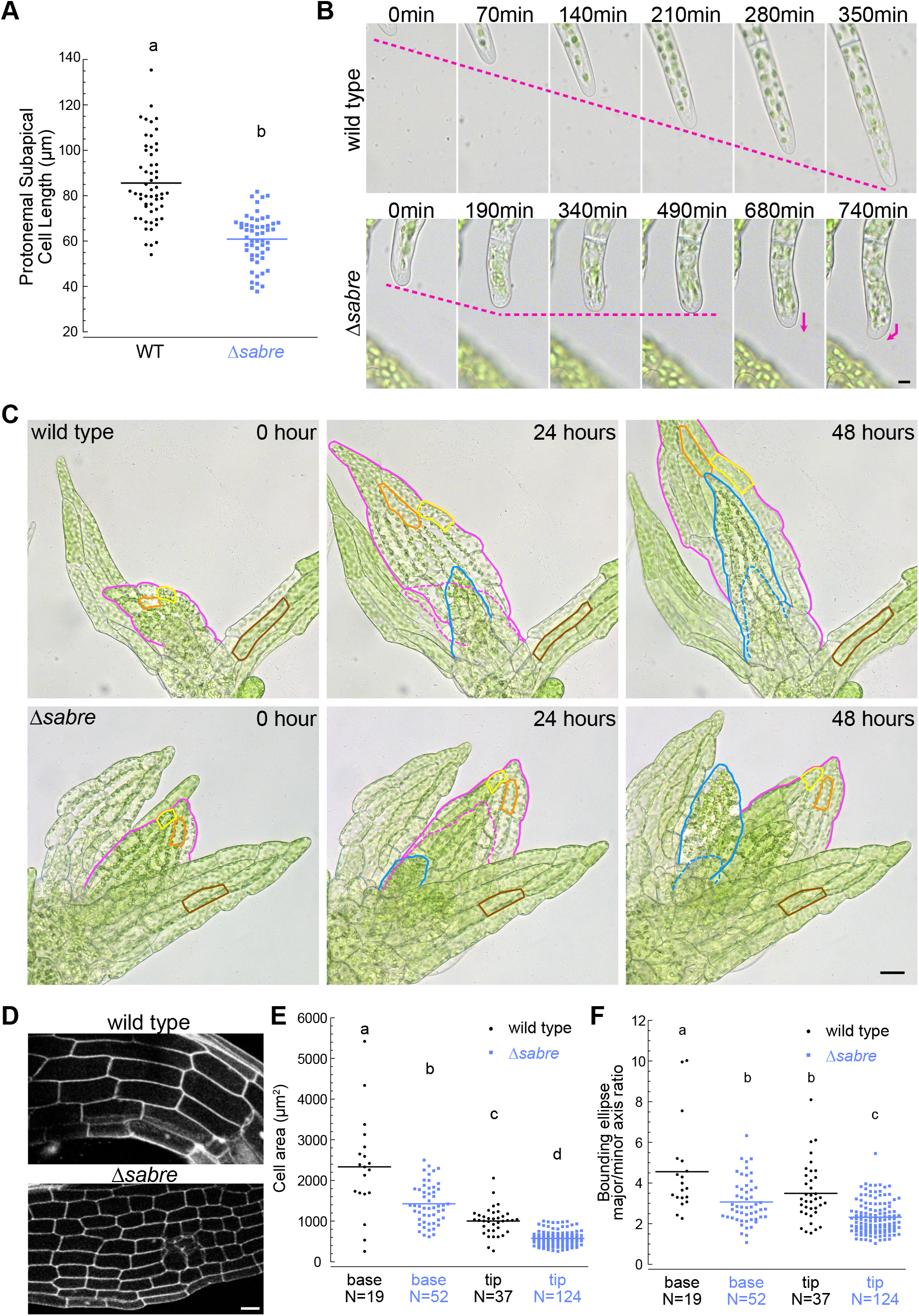
Reduced cell expansion underlies small plant size in *Δsabre*. (A) Quantification of sub-apical cell length from 7 -day-old plants regenerated from protoplasts. N=56, wild type; N=53, *Δsabre*. Letters indicate groups with significantly different means as determined by a Student t-test for unpaired data with equal variance, p<0.0001. (B) Bright field time-lapse images for wild type (top) and *Δsabre* protonemata (bottom). Scale bar, 5μm. Magenta dashed lines indicate the apical positions of the cells. Magenta arrows indicate growth directionality. Also see Video 2. (C) Bright field time-lapse imaging of gametophores growing in microfluidic imaging devices. Magenta and blue lines indicate example phyllids 1 and 2, respectively, that expanded during the imaging period. Dashed lines in corresponding colors indicate the phyllid outlined 24 hours before. Orange and yellow lines outline example cells that expanded during the imaging period. Brown lines highlight example cells in a mature phyllid that did not obviously increase in size. Scale bar, 20 μm. Also see Video 3. (D) Example confocal fluorescent images of phyllids stained with propidium iodide (PI) used for quantification in (E-F). Scale bar, 30 μm. (E) Quantification of phyllid cell area. Base and top indicate cells were located at the base or top of the phyllid. (F) Quantification of the ratio of major/minor axis of the bounding ellipse fitted to each cell. Number of cells in each category indicated under the graph. Letters indicate groups that are significantly different as determined by one-way ANOVA with Tukey’s HSD post-hoc test (α=0.05).

Interestingly, *Δsabre* protonemal cells often changed direction during growth (Figure 2B – video 2). Both actin and microtubules contribute to maintaining the direction of polarized growth. Cytoplasmic microtubules polymerize towards the cell tip where their plus ends then focus onto an apically localized actin spot (Hiwatashi et al., 2014; Wu et al., 2018). When actin filaments are disrupted, microtubules no longer focus below the tip (Wu and Bezanilla, 2018) and growth is inhibited. When microtubules are disrupted, the actin spot randomly appears and disappears throughout the cell, with expansion occurring in areas that accumulate actin, ultimately resulting in a loss of directional growth (Wu and Bezanilla, 2018; Yamada and Goshima, 2018). Since, defects in growth directionality in *Δsabre* cells resembled wild type cells lacking microtubules (Doonan et al., 1988; Wu and Bezanilla, 2018), we wondered if SABRE influenced the microtubule focus at the cell apex. However, similar to wild type, in *Δsabre* cells the actin and microtubule foci were persistently present at the tip (Figure 2 – figure supplement 1A, B). Thus, the observed defects in growth directionality and rate appear to be independent of actin and microtubules. Actin and microtubules also form dynamic networks along the cell cortex of protonemal cells. In Arabidopsis, cortical microtubule orientation was altered in *sabre* mutants from mostly transverse to random in root epidermal cells (Pietra et al., 2013). In contrast to the observations in Arabidopsis, we did not observe obvious changes in microtubule organization or dynamics at the cortex (Figure 2 – figure supplement 1C). Interestingly, cortical actin labeled with Lifeact-GFP exhibited a slight increase in dynamics (Figure 2 – figure supplement 1D), suggesting that in *P. patens SABRE* may impact actin instead of microtubules.

Diffuse growing tissues in the *P. patens Δsabre* mutants were stunted similar to what was observed in the Arabidopsis *sabre* mutant. Time-lapse imaging of expanding phyllids (Figure 2 – video 3) revealed that due to defective cell expansion (Figure 2C yellow and orange lines), *Δsabre* grew significantly less within the same time window compared to wild type (Figure 2C, magenta and blue lines and dashes). In mature phyllids, the final cell size was also smaller in *Δsabre* (Figure 2C brown lines). We quantified cell size in fully expanded phyllids and discovered that cell area at the base and tip of phyllids was reduced in *Δsabre* (Figure 2D-F), consistent with the overall stunted gametophore stature (Figure 1D-F). In particular, cell area at the base of phyllids, which are generally the largest cells in the phyllid, were more affected than cells at the tip of the phyllid in *Δsabre* (Figure 2E), suggesting that in the absence of SABRE function cells may not be able to expand beyond a certain size. To measure changes in shapes of individual cells, we fit a bounding ellipse to each cell, and quantified the ratio between the major and minor axes. Cells were less elongated in *Δsabre* both at the tip and the base of the phyllids as compared to wild type (Figure 2F).

### Loss of SABRE results in cytokinetic delays and can lead to failures in cytokinesis

During cytokinesis plant cells use the phragmoplast, a microtubule-based structure, to build a new cell plate that physically separates the daughter cells. In protonemata, the phragmoplast forms between the two daughter nuclei, expanding perpendicular to the axis of the filament and eventually fusing with the plasma membrane. Phragmoplast microtubules – a dense bipolar array of microtubules direct late secretory vesicles to the midzone where they fuse to form a membrane encapsulated cell plate (Smertenko et al., 2018). During the early stage known as the disc phragmoplast, interdigitated microtubules with their plus ends at the cell equator are arranged in a spindle-like structure to establish the phragmoplast. As the cell plate grows, the microtubules label the outer edge of the expanding cell plate, known as the ring phragmoplast. Eventually the developing cell plate reaches the existing side wall and fuses with it, at which point the microtubule array dissipates (Boruc and Van Damme, 2015; Smertenko, 2018). In *Δsabre* protonemata, we discovered that while mitosis is unaltered (for example, Figure 3A 0-12 min), phragmoplast microtubules were present for significantly longer than in control cells (Figure 3A, figure 3 – video 4). We found that starting at the time of disc phragmoplast establishment (Figure 3A 12 min), all control cell phragmoplast microtubules disappeared within 28 to 50 minutes (N=10). However, for all *Δsabre* cells (N=10), microtubules were still present for at least 50 minutes after disc phragmoplast establishment. To test whether the delay in microtubule disassembly observed in *Δsabre* phragmoplasts resulted from defects in microtubule dynamics, we performed photobleaching on phragmoplast microtubules labeled with GFP-tubulin (Figure 3 – figure supplement 1A). Microtubule fluorescence recovered from photobleaching similarly in control and *Δsabre* cells (Figure 3 –figure supplement 1B), indicating that *SABRE* does not regulate microtubule dynamics during phragmoplast expansion.

**Figure 3.**
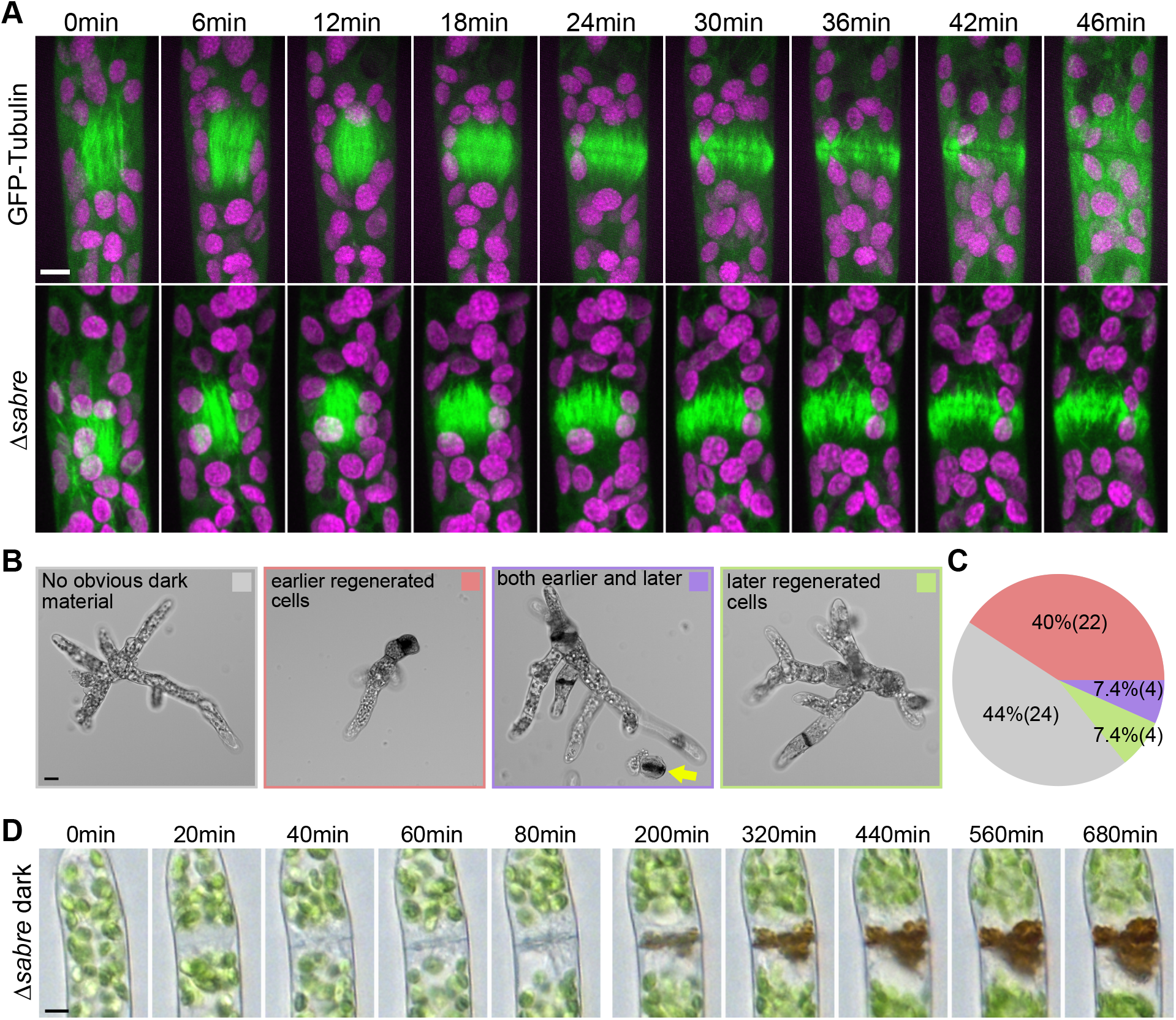
Delays in disassembling phragmoplast microtubules and quantification of cytokinesis failures during protonemal development. (A) Time-lapse imaging of phragmoplast microtubules. GFP-tubulin (green) and chlorophyll autofluorescence (magenta) are shown. First frames (0 min) occur within 2 min of nuclear envelope breakdown. Scale bar, 5 μm. Also see Video 4. (B) Representative images depicting brown material deposition near cell plates in 7-day old plants regenerated from protoplasts. Colored frames correspond to categories quantified in C. Yellow arrow indicates an example of a dead protoplast containing dark brown material at the first cell division site resulting in the failure to regenerate. Scale bar, 20 μm. (C) Frequency of brown material deposits at different developmental stages (marked by cell shape and position in regenerated plant). Numbers in parentheses indicate numbers of plants. (D) Bright field time-lapse EDF images of brown material deposition in a cell that underwent cell division. Also see Video 5. Scale bar, 5 μm.

Besides the lengthy delay in phragmoplast microtubule disassembly in protonemata, we observed that a fraction of cells contained dark brown material, which accumulated near the cell division plane (Figure 3B-D). We used numerous dyes for cell wall components to attempt to stain the brown material to get a hint of its composition, but none of these dyes stained. Time-lapse imaging revealed that brown material deposition was slow (Figure 3D, figure 3 – video 5), but always initiated during a cell division event. Cells with brown material would either after a long time recover and grow or die (figure 3 – video 5). Cell death suggests that defects in cell plate formation resulted in loss of cell integrity. Interestingly, brown material appeared more frequently in cells with a large diameter, characteristic of the first few cells in plants regenerated from protoplasts (Figure 3B, C). Cells with a large diameter have inherently more degrees of freedom for orienting the phragmoplast. Furthermore, phragmoplast expansion and insertion occurs over a longer distance in these cells. We also noticed that *Δsabre* protoplasts regenerated inefficiently compared to wild type, likely because many protoplasts died during the first cell division, with brown material deposited at the cell division site (Figure 3B yellow arrow). Considering that neither cell length nor average growth rate accounted for the 60% reduction in protonemal area (Figure 1, 2), we reasoned that the developmental delay caused by delays and failures in cytokinesis coupled with frequent pauses in growth likely account for the rest of the decrease in protonemal plant size.

### SABRE co-localized with a fraction of the ER at the cell tip, cell cortex and in the phragmoplast midzone during cell plate maturation

To determine how *SABRE* impacts cell growth and division, we generated fluorescent fusions of *SABRE* to analyze its subcellular distribution. We inserted sequences encoding for three tandem mEGFP proteins (Vidali et al., 2009) at the native locus of *SABRE* using CRISPR-Cas9 mediated homology directed repair (HDR). However, the SABRE-3XmEGFP signal was extremely weak. To increase the signal, we generated a C-terminal tag with three tandem mNeonGreen proteins (hereafter, SAB-3mNG) (Figure 4 – figure supplement 1A). We demonstrated that tagging SABRE at its C terminus did not influence its function, as we observed no growth defects in tagged plants (Figure 4 – figure supplement 2A).

Confocal microscopy revealed that SABRE-3mNG formed small puncta at the cell cortex, whose density was higher near the tip of the apical cell (Figure 4A). To further increase the SABRE signal, we inserted the maize ubiquitin promoter (a strong promoter) before the start codon of *SABRE-3mNG* coding sequence at the native genomic locus (Figure 4 – figure supplement 1B). We found that OE-SAB-3mNG plants were indistinguishable from SAB-3mNG plants, demonstrating that overexpression did not have any adverse consequences for protonemal growth (Figure 4 – figure supplement 2B). Other than a higher density of SABRE puncta, we observed a similar localization pattern in the overexpression line (Figure 4 – figure supplement 2C). Because SABRE-3mNG was challenging to detect as the focal plane moved deeper into the cell, we used OE-SAB-3mNG for confocal imaging during cell division. We found that OE-SAB-3mNG formed discrete puncta decorating the entire developing cell plate (Figure 4B). Note that due to weak signals in both SABRE-3mNG and OE-SAB-3mNG, autofluorescence from the chloroplasts was prominently visible in the mNeonGreen channel for both lines.

**Figure 4.**
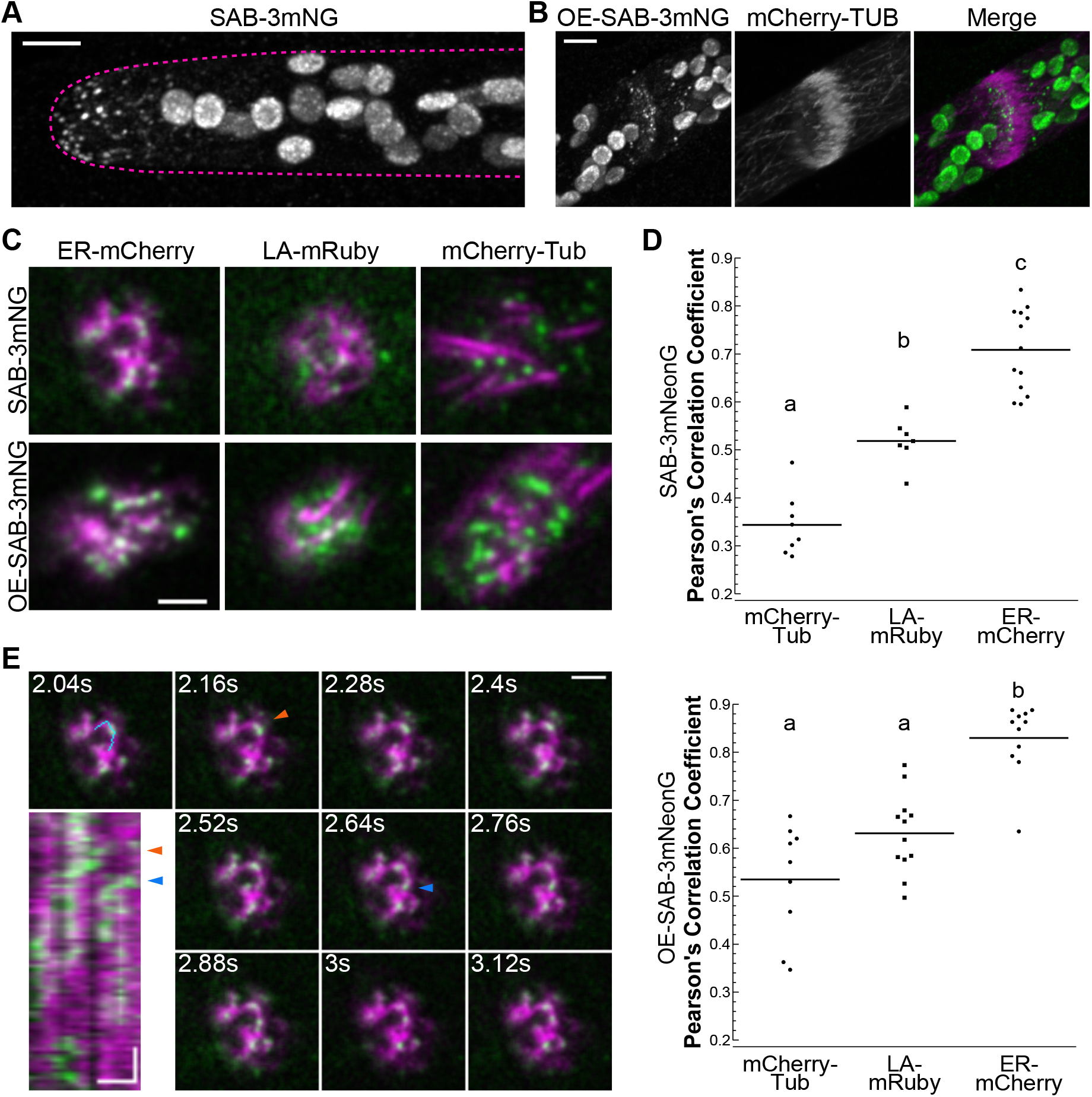
Localization of SABRE revealed by tagging SABRE at the C-terminus with three tandem mNeonGreen proteins. (A) SABRE-3mNeonGreen (SAB-3mNG) forms small puncta at the cell cortex that are more numerous near the tip of a growing apical cell. Image is a maximum projection of a deconvolved confocal z-stack. Magenta dashed line indicates the outline of the cell. Scale bar, 5 μm. (B) Deconvolved confocal images of SABRE localization at the maturing cell plate. Scale bar, 5 μm. SAB-3mNG (green in merge) and mCherry-tubulin (magenta in merge) are shown. Large globular structures are chloroplasts, which autofluorescence in the mNeonGreen channel under these imaging conditions. (C) VAEM images of SAB-3mNG (green) with mCherry-tubulin (mCherry-Tub), lifeact-mRuby (LA-mRuby) and mCherry-KDEL (ER-mCherry) (magenta). Representative images are the first frame of a VAEM time-lapse acquisition. Scale bar, 2 μm. Also see Video 6. (D) Quantification of SAB-3mNG (top) and OE-SAB-3mNG (bottom) co-localization with markers shown in C. Pearson’s correlation coefficient from the first 10 frames of a VAEM time-lapse acquisition of a cell taken every 120 ms was calculated, the average correlation coefficient for each cell is plotted. For SAB-3mNG: N=8, mCherry-Tub; N=7, LA-mRuby; N=13, ER-mCherry. For OE-SAB-3mNG: N=9, mCherry-Tub; N=12, LA-mRuby; N=11, ER-mCherry. Letters indicate groups with significantly different means as determined by ANOVA with a Tukey’s HSD all pair comparison post-hoc test (α=0.05). (E) VAEM time-lapse acquisition showing SAB-3mNG (green) moving along an ER tubule (magenta). Scale bar, 2 μm. Kymograph was generated using the cyan line drawn in the 2.04 sec frame. Orange and blue arrowheads indicate 2 events when a SAB-3mNG puncta starts to move along the ER tubule. Horizontal scale bar, 1 μm. Vertical scale bar, 1.2 seconds.

During cell division, microtubules form the phragmoplast, actin interacts with microtubules and guides the expanding phragmoplast (Buschmann and Müller, 2019; Müller, 2019; Wu and Bezanilla, 2014), while the endoplasmic reticulum (ER) threads through the developing cell plate to build plasmodesmata – plant-specific channels that connect adjacent plant cells (Sager and Lee, 2018; Tilney et al., 1991). Since *Δsabre* plants have severe defects in protonemal cell division, we generated SAB-3mNG and OE-SAB-3mNG in moss lines possessing microtubules (mCherry-tubulin), actin (Lifeact-mRuby) and endoplasmic reticulum (ER luminal marker SP-mCherry-KDEL) markers, enabling inquiry of SABRE behavior in the context of known phragmoplast structures. Initially, to maximize the SABRE signal, we used variable angle epifluorescence microscopy (VAEM) to image SABRE simultaneously with these markers at the cell cortex. Consistent with the finding that microtubules were not affected in *Δsabre* plants (Figure 2 – figure supplement 1A, C) and that SABRE localizes to the nascent cell plate even in late phragmoplasts that lack microtubules (Figure 4B), we found that dynamic cortical SABRE puncta did not associate with microtubules (Figure 4C, figure 4 – video 6). We did observe limited SABRE interactions with actin at the cell cortex (Figure 4C, figure 5 – video 6), which could explain our earlier finding of increased actin dynamics in *Δsabre* (Figure 2 – figure supplement 1D). However most surprisingly, we discovered that cortical SABRE co-localized with the ER (Figure 4C). Measuring Pearson’s correlation coefficient demonstrated strong spatial correlation between ER and SABRE, but not between ER and both microtubules and actin (Figure 4D, top). We also observed spatial and temporal correlation between ER and SABRE, with SABRE puncta moving along ER tubules (Figure 4E, video 6). In a kymograph along one tubule highlighted by the blue line, there were two events where SAB-3mNG dots translocated along the tubule (Figure 4E arrowheads). Overexpression of SABRE did not dramatically alter SABRE localization at the cell cortex other than to uniformly increase the value of the correlation coefficients likely resulting from increased number of SABRE puncta in the imaging field (Figure 4D, bottom).

**Figure 5.**
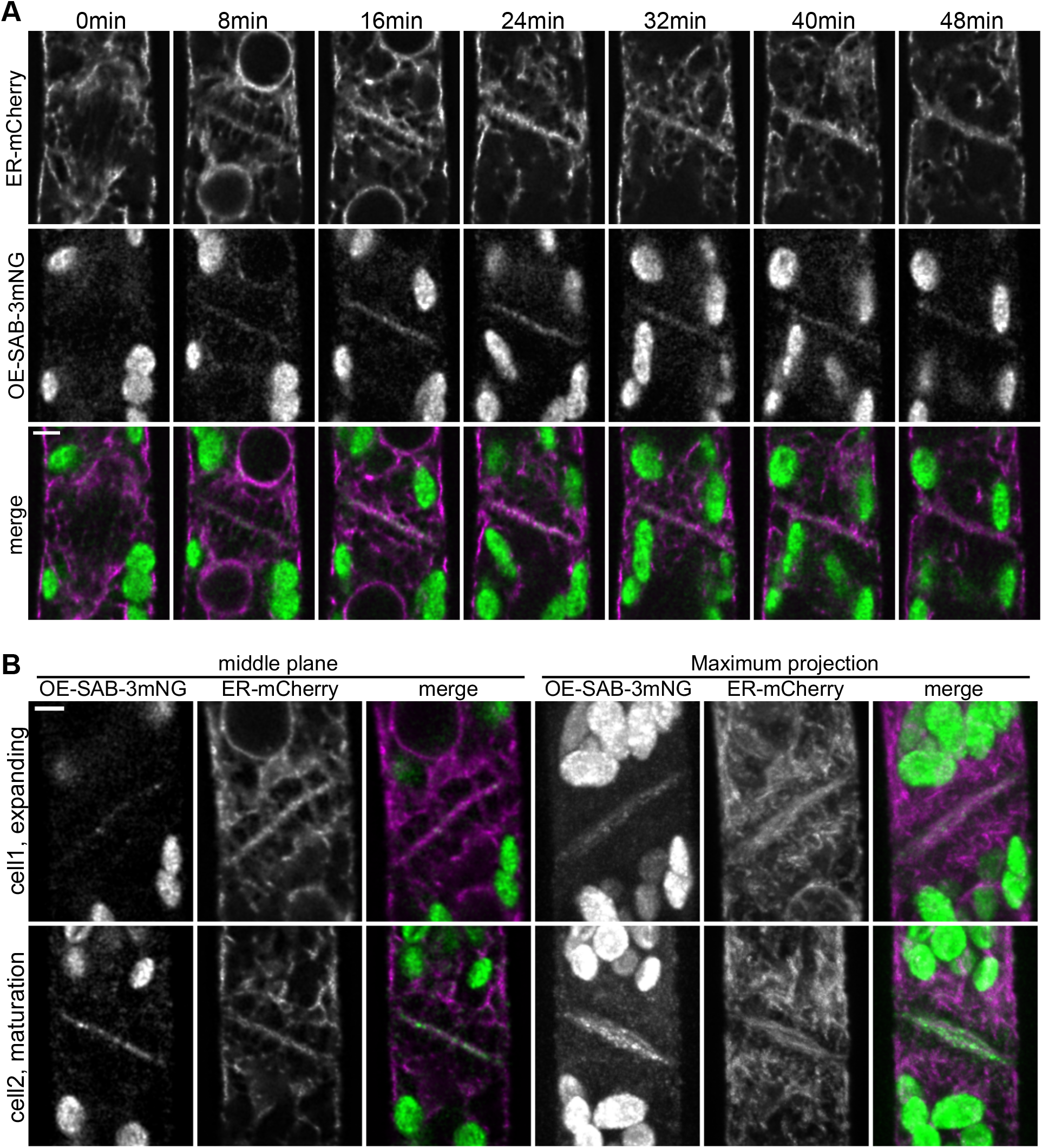
SABRE and ER accumulate on the nascent cell plate during cell plate maturation. (A) Deconvolved confocal images of OE-SAB-3mNG (green in merge) and mCherry-tubulin (magenta in merge) during cell division. Individual frames are medial focal planes of the cell. Scale bar, 3μm. Also see Video 7. (B) Two representative cells with ER-mCherry (magenta in merge) and OE-SAB-3mNG (green in merge) showing the difference in the accumulation of SABRE and ER at during cell plate expansion and maturation. Scale bar, 3 μm.

Since *Δsabre* exhibited serious defects in cell plate maturation and co-localized with the ER at the cell cortex, we further investigated the timing of SABRE and ER localization during cell division. After mitosis, the disc phragmoplast was labeled with thin strands of ER parallel to the microtubules and very little ER was present in the phragmoplast midzone (Figure 5 – figure supplement 1A first timepoint). In the ring phragmoplast, microtubules were shorter and expanded with the phragmoplast edge (Figure 5 – figure supplement 1), while the ER strands parallel to the microtubules became more defined and the ER accumulated in the midzone perpendicular to the microtubules lining the future cell plate (Figure 5 – figure supplement 1A). Cell plate maturation occurs during the late phragmoplast stage (Smertenko et al., 2017), coincident with an increase in the ER signal (3^rd^ and later timepoints of Figure 5A, figure 5 – figure supplement 1, Figure 5B). Confocal time-lapse imaging revealed that OE-SAB-3mNG localized to the midzone of the ring phragmoplast weakly, and the signal strengthened as the phragmoplast fully expanded and inserted into the side wall (Figure 5A, B, figure 5 – figure supplement 1B). The strongest OE-SAB-3mNG signal correlated with the timing of cell plate maturation (Figure 5A, B, video 7) and remained until the ER signal visibly split in two on either side of the new cell plate (last 2 time points of Figure 5A).

### Loss of SABRE impacts ER morphology in the cytoplasm during interphase and at the cell plate during cell division

Given the striking co-localization between SABRE and the ER, we further analyzed ER morphology and dynamics in *Δsabre*. With an ER luminal marker, SP-GFP-KDEL, we were able to examine the overall ER structure in the cytoplasm. We discovered that *Δsabre* cells contained abnormal ER aggregates in the cytoplasm and at the cell cortex in protonemata, which might underlie the defects in polarized growth rate and directionality (Figure 6A, video 8). Initially during cell division, ER localization was normal (0 min timepoint of Figure 6B, video 9). However, during the transition to an increase in ER parallel to the cell plate, the cell plate ER signal noticeably buckled in *Δsabre*, while in control cells the ER was straight (Figure 6B magenta arrowheads, video 9). Interestingly buckling occurred when SABRE maximally accumulated on the cell plate in control cells (Figure 5). Note that while twisting of the ER was obvious in the center of the cell plate, the edges adjacent to the side wall remained fixed, indicating that phragmoplast guidance mechanisms and the phragmoplast insertion site were not affected (Figure 6B, figure 6 – video 9). For *Δsabre* cells that divided relatively normally, the twisted ER, which was observed in all *Δsabre* cells (N=31), eventually straightened out 20 to 30 minutes later (Figure 6B, figure 6 – video 9).

**Figure 6.**
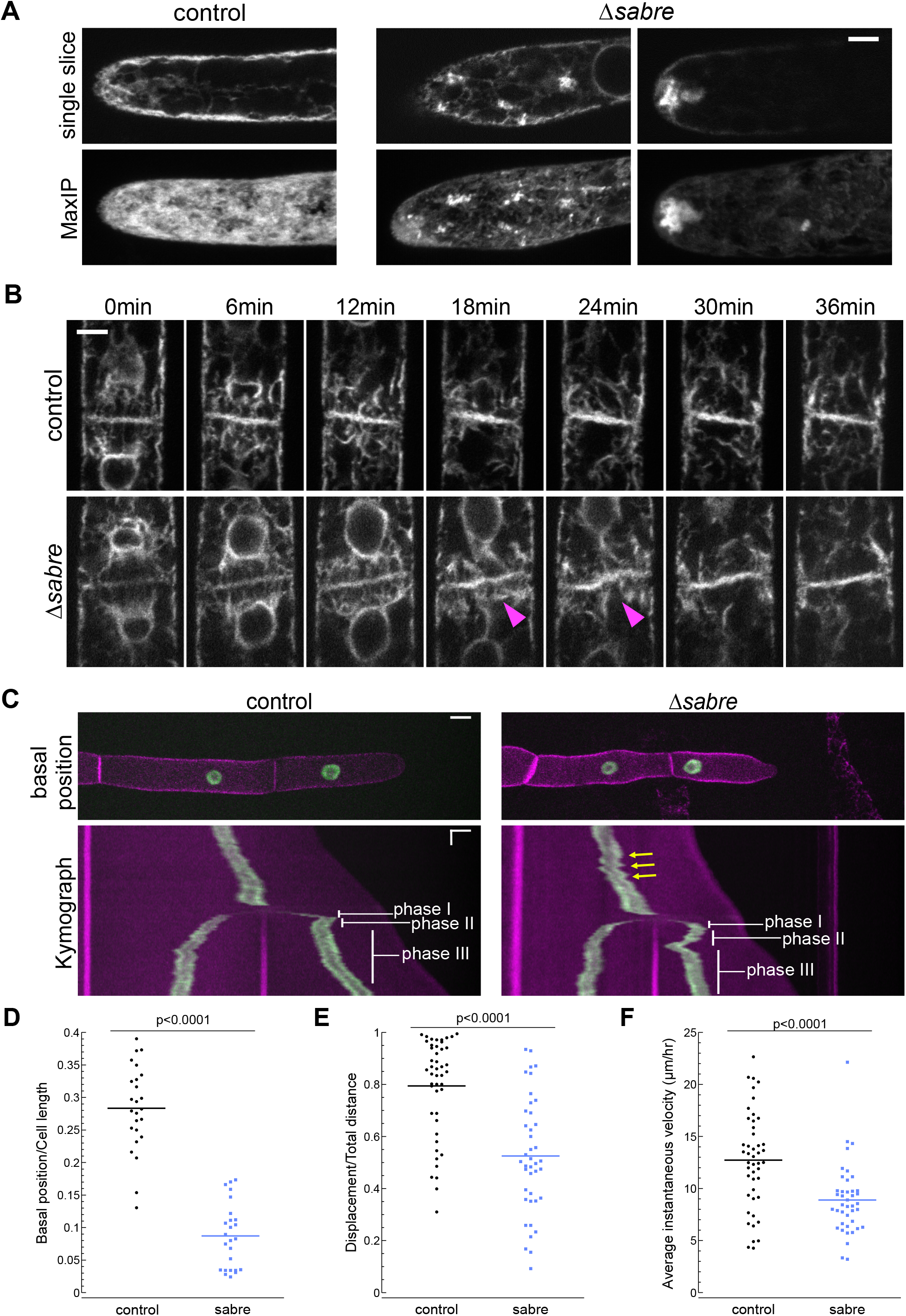
SABRE influences ER morphology and nuclear migration. (A) ER labeled with SP-GFP-KDEL in the cytoplasm of growing apical cells. Top: single focal plane. Bottom: maximum projection of confocal Z-stacks. Scale bar, 5 μm. Also see Video 8. (B) ER during cell division in wild type and *Δsabre* cells. First frame is within 3 minutes of nuclear envelop reformation. ER buckles at the nascent cell plate in *Δsabre* (magenta arrowheads). Scale bar, 5 μm. Also see Video 9. (C) Top panel: representative images of control and *Δsabre* cells when the nucleus in the tip cell is closest to the new cell plate (basal position). NLS-GFP-GUS (green) accumulates in the nucleus, and SNAP-TM-mCherry (magenta) labels the plasma membrane. Scale bar, 10 μm. Bottom panel: Kymograph created by drawing a line in the middle of the cell along the growth axis. Horizontal scale bar, 10 μm. Vertical scale bar, 1 hour. (D) Distance of the nucleus in the apical cell to the newly formed cell plate when it is at the basal position normalized to cell length. N = 25. (E) Quantification of linearity of nuclear movement during apical moving phases (I and III). Linearity determined by the ratio of total distance traveled divided by total linear displacement. Nuclear movement was tracked with TrackMate plugin in Fiji generating distance and displacement of nucleus. N=46, control; N=41, *Δsabre*. (F) Quantification of nuclear velocity for the same cells and migration phases in E. Average instantaneous velocity (μm/hour) was determined by dividing the total distance traveled by the time. Statistical analyses for D-F were performed using Student t-test for unpaired data with equal variance. Also see Video 11.

In addition to ER buckling, we noticed that nuclei in *Δsabre* apical cells exhibited aberrant motility after cell division, as ER also outlines the nuclear envelope. Nuclei in control cells (Figure 6 – figure supplement 1A), moved apically (phase I, blue arrows, see also Figure 6C), basally (phase II, orange arrows, see also Figure 6C) and then resumed apical migration (phase III, white arrows, see also Figure 6C) as has been described previously (Yamada and Goshima, 2018). In *Δsabre* cells, the initial apical movement of the nucleus was delayed in 39% of the 31 imaged cells (Figure 6 – figure supplement 1B green arrowheads – video 10), resulting in a close association between the nucleus and the cell plate, and coinciding with ER buckling (Figure 9 - supplement magenta arrowheads – video 10). While the phase I apical nuclear movement was normal in the remaining 61% of imaged *Δsabre* cells, the subsequent basal movement was dramatically exaggerated in all cells (Figure 6 – figure supplement 1 red arrowheads – video 10).

To further quantify nuclear migration defects, we disrupted *SABRE* in a line expressing a nuclear localized GFP (NLS-GFP-GUS) and a plasma membrane marker (SNAP-TM-mCherry) (van Gisbergen et al., 2018, p. 10), enabling imaging of cell division and nuclear movement before and after cell division (Figure 6C – video 11). In control cells, basal migration (phase II) was often subtle or sometimes even appeared to be a stationary phase (Figure 6C) (Yamada and Goshima, 2018). In contrast, *Δsabre* basal (phase II) nuclear movement was extreme (Figure 6C – video 11). In many cases, the nucleus migrated so far back that it appeared to deform as it smashed up against the new cell plate (Figure 6C – video 11). To quantify the defect in basal movement, we measured the distance between the nucleus and the cell plate when the nucleus was closest to the cell plate (basal position), normalizing the basal position to the apical cell length at that timepoint. Despite shorter cells in *Δsabre*, the relative basal nuclear position was significantly smaller than in control cells (Figure 6D). Beyond the immediate nuclear migration defects after cytokinesis, we also observed less consistent velocity and directionality during interphase in *Δsabre*, represented by the zigzagging trajectory in the kymograph (Figure 6C – yellow arrows, video 11). We tracked the nucleus and quantified the ratio between the final nuclear displacement and the total distance traveled in interphase. We discovered that *Δsabre* nuclear movement was less linear and the ratio was smaller (Figure 6E) as compared to wild type. The average instantaneous velocity of the nucleus in *Δsabre* was expectedly smaller, since the cells were not always actively growing (Figure 6F). Given that SABRE influences the ER and that the nuclear envelope is contiguous with the ER, defects in nuclear migration likely result from altered ER function and suggest that SABRE may provide a link between the nuclear envelope and motors responsible for nuclear migration (Leong et al., 2020; Yamada and Goshima, 2018).

To characterize key components of the cell division machinery in *Δsabre* cells that failed to form a normal cell plate and resulted in cell death, we imaged the ER, lifeact-GFP labeling actin and GFP-tubulin labeling microtubules. We observed the ER signal transition from defined tubules (Figure 7A, 40min) in the phragmoplast to a diffuse signal (Figure 7A, 50 min), correlating with the onset of division failure. As the brown material accumulated, it was outlined by the ER signal (Figure 7A, figure 7 – video 12). Both lifeact-GFP and GFP-tubulin signals were relatively normal up until the phragmoplast had expanded to the parental cell membrane. However, at that point lifeact-GFP became uneven around the phragmoplast with large gaps appearing near the presumptive cell plate. Lifeact-GFP lingered near the cell plate as the brown material generated a gap in the fluorescence gradually “invading” the cytoplasm, which was accompanied by flashes of lifeact-GFP fluorescence (Figure 7B, 50 min and after). In young plants regenerating from protoplasts, phragmoplasts labeled with GFP-tubulin expand across a larger distance, often finishing insertion on one side of the cell and then extending to the other (Figure 7C). In *Δsabre* cells that accumulated brown material, phragmoplast microtubules on the outer edge became disoriented once they reached the parental plasma membrane (Figure 7C, 160 min). Later, deposition of brown material formed a gap between the remaining phragmoplast microtubules (Figure 7C). Sometimes cell plate defects resulted in immediate death. These cells lysed quickly after division and exhibited similar trends to cells that accumulated brown material; phragmoplast expansion was normal until it reached the side wall, at which point the cell lysed (Figure 7 – figure supplement 1). Taken together our data suggest that phragmoplast expansion is normal in *Δsabre*. However, during cell plate maturation, defects arise in *Δsabre* cells; the ER becomes diffuse and both the actin and microtubule cytoskeletons remain associated with the cell plate.

**Figure 7.**
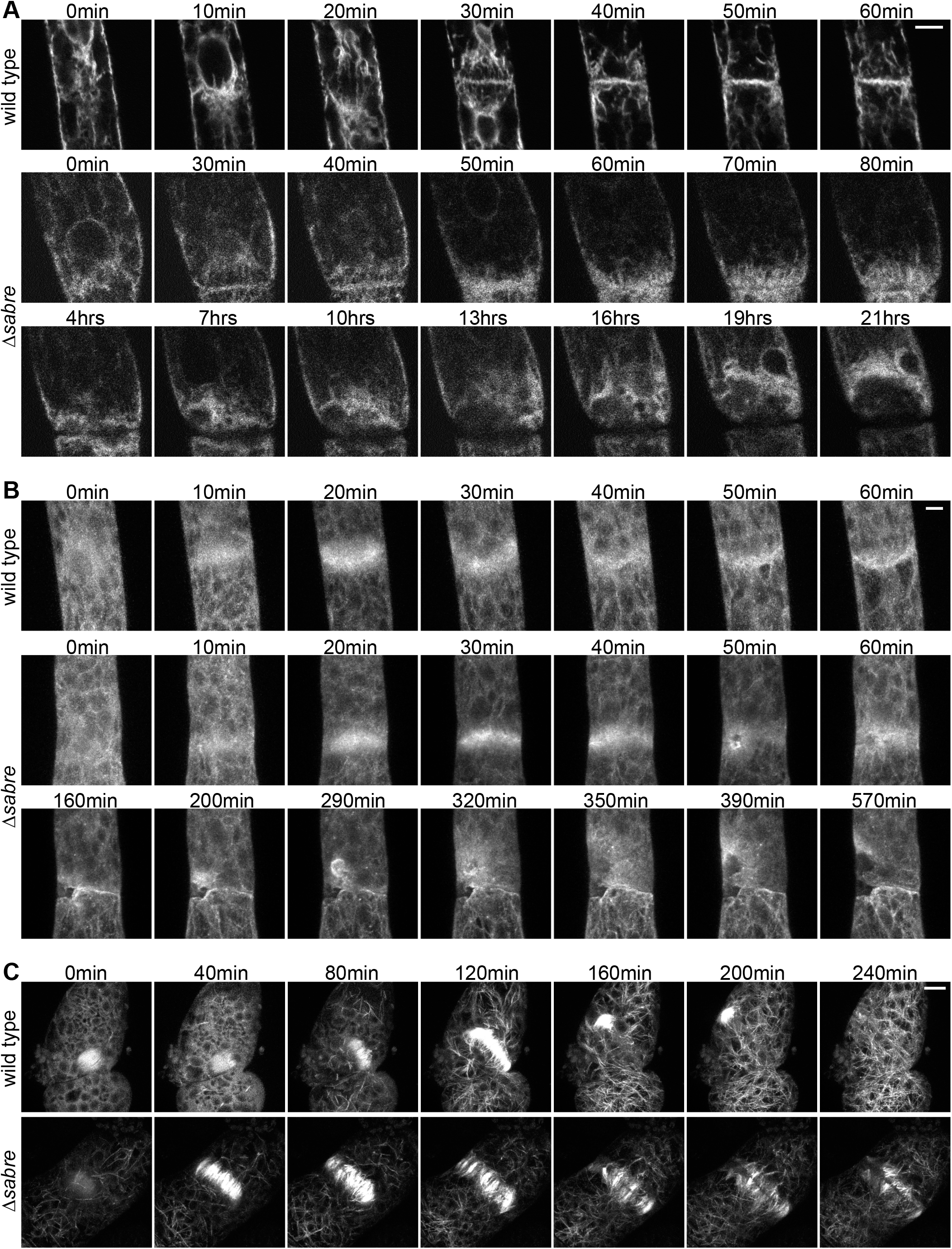
Behavior of ER, actin and microtubules during cytokinesis failures in *Δsabre*. Single focal planes from confocal images of (A) ER labeled with SP-GFP-KDEL, maximum projection of confocal Z-stacks of (B) actin labeled with Lifeact-GFP, and (C) microtubules labeled with GFP-tubulin in wild type and *Δsabre* cells that accumulate brown material during cell division. (A) Between 0-20 min a phragmoplast formed. At 30 min, the *Δsabre* phragmoplast failed to mature and brown material deposition gradually occurred over the following hours. Scale bar, 6 μm. (B) Scale bar, 5 μm. (C) Scale bar, 10 μm. Also see Video 12.

### SABRE impacts callose deposition

To follow membrane and cell wall remodeling events occurring during cell division, we imaged cell division in the presence of the lipophilic dye, FM4-64, and the callose-specific dye, aniline blue. FM4-64 labels endocytic membranes (Aniento and Robinson, 2005; Jelínková et al., 2010; Klima and Foissner, 2008; Kutsuna and Hasezawa, 2002; Tse et al., 2004; Ueda et al., 2004; van Gisbergen et al., 2008), which are readily incorporated into the membrane surrounding the nascent cell plate early in cytokinesis as clathrin-mediated endocytosis is integral for remodeling the tubular membrane network during early phragmoplast formation (Dhonukshe et al., 2006; Lam et al., 2008; Zhang et al., 2011). However, once the phragmoplast has expanded across the entire mother cell, FM4-64 labeling decreases, consistent with a transition to different membrane trafficking machinery employed specifically during cell plate maturation and membrane fusion (Drakakaki, 2015; Park et al., 2014; Smertenko et al., 2017). In seed plants, callose is deposited early accumulating during the ring stage of the phragmoplast and reaching a peak just prior to fusion of the membrane encapsulating the nascent cell plate with the parental plasma membrane (Samuels et al., 1995). By imaging actively dividing wild type cells, we observed that aniline blue fluorescence steadily increased following a characteristic decrease in FM4-64 staining (Figure 8A, figure 8 – video 13). Notably, aniline blue only stained cell plates that had fully expanded, suggesting that during phragmoplast expansion, callose within the membranous tubular network is not accessible to aniline blue in the extracellular environment. However, as FM4-64 levels diminished occurring during cell plate maturation and fusion of the cell plate membrane with the parental plasma membrane (Figure 8 – figure supplement 1), aniline blue could access the callose leading to the observed increase in staining (Figure 8A). *Δsabre* cells that divided relatively normally (contained no brown material) exhibited a similar decrease in FM4-64 intensity (Figure 8B solid lines). However, the FM4-64 signal continued to decrease for an additional ten minutes, before aniline blue began to rise. At this time, wild type cells would have been fully stained with aniline blue. Further the rate of increase in aniline blue was significantly slower in *Δsabre* cells (Figure 4B, dashed lines), suggesting that it either took longer for aniline blue to diffuse into the cell plate or that callose was deposited later in *Δsabre* cells.

**Figure 8.**
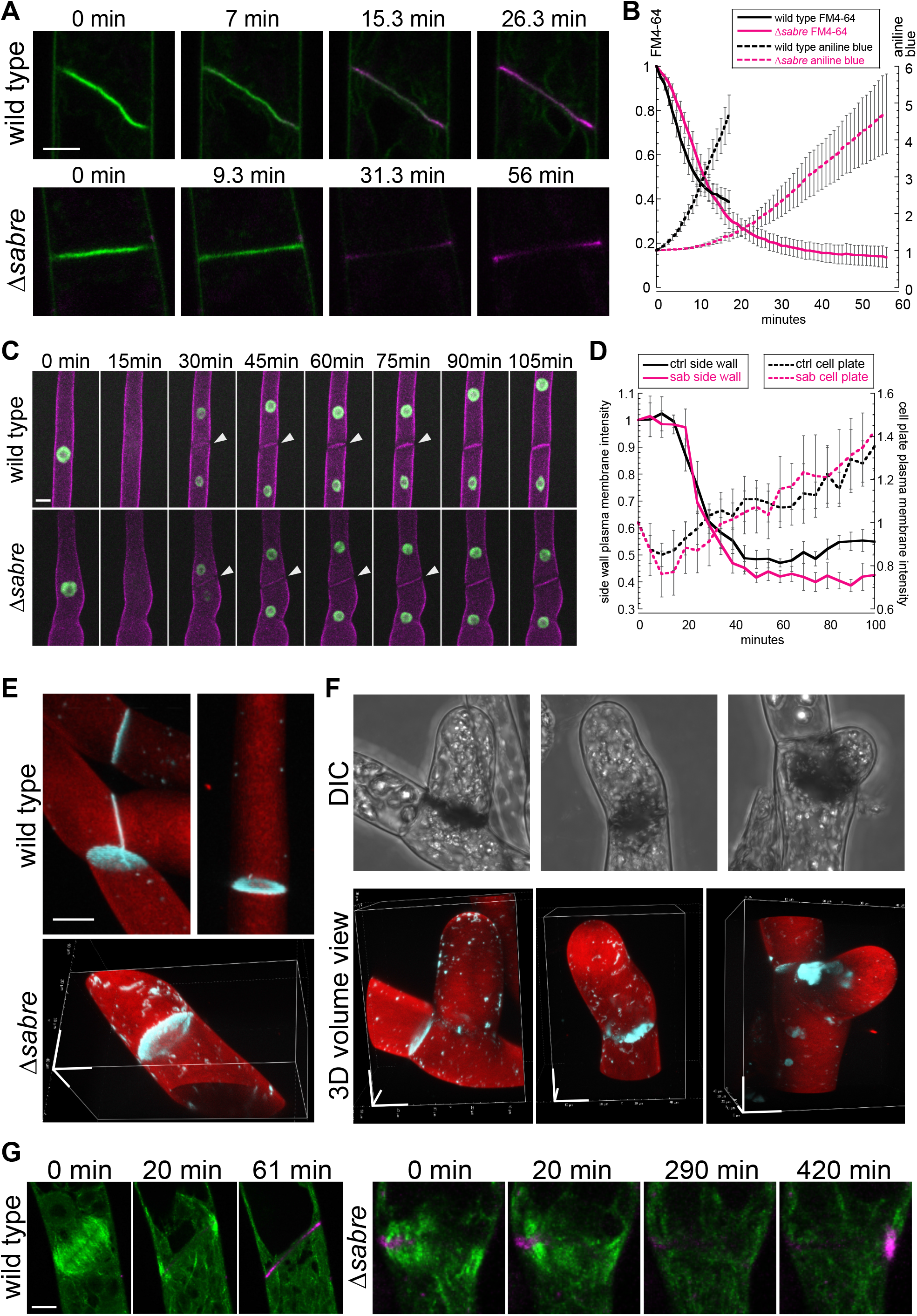
Aniline blue deposition is altered in *Δsabre*. (A) Time lapse of FM4-64 (green) and aniline blue (magenta) showing aniline blue staining the cell plate after FM4-64 staining diminishes. Also see Video 13. (B) Quantification of FM4-64 and aniline blue at the developing cell plate over time. Intensity is measured by drawing a 15-pixel wide curved line along the developing cell plate and measuring the mean intensity value. Each data point is the average of N = 6 cells for each category. Intensity is normalized to the starting point of the time lapse – the moment when FM4-64 is the strongest and no aniline blue stain accumulates. Error bars, standard error of the mean. (C) Time lapse of membrane marker SNAP-TM-mCherry (magenta) and nuclear marker NLS-GFP-GUS (green) during cell division. 0-minute frame depicts the last time point (within 5 minutes) before nuclear envelope break down. Also see Video 11. (D) Quantification of signal intensity in (C). A region of interest of 6-10 micron^2^ was manually drawn at the side wall where membrane fusion occurs and at the middle of the cell plate respectively, mean intensity was measured for those two regions of interests and plotted against time. Each data point is the average of N = 6 cells for each category. Error bars, standard error of the mean. (E) Example 3D images of normal mature cell plates stained with aniline blue (cyan) and fast scarlet (F2B) (red) to outline the cell wall. (F) Examples of different aniline blue staining patterns in *Δsabre* mutants containing brown material. Left, weak stain. Middle, partial stain. Right, large chunk near the cell plate. Scale bars, 10 μm. (G) Time lapse showing aniline blue staining (magenta) in a WT cell and a *Δsabre* cell with brown material. Images are maximum projections of deconvolved confocal z-stacks. Microtubules labeled with GFP-tubulin are shown in green. Scale bar, 5 μm. Also see Video 14.

To distinguish between defects in membrane fusion or delays in callose deposition, we analyzed images of cell division in protonemata labeled with a nuclear GFP and the membrane marker SNAP-TM-mCherry (Figure 6C). In these lines, loss of GFP fluorescence from the nucleus signaled nuclear envelope break down and the onset of prometaphase. In contrast to FM4-64, SNAP-TM-mCherry did not accumulate on the nascent cell plate (Figure 8C, figure 6 – video 11). Instead, SNAP-TM-mCherry appeared at the cell plate 30 min after nuclear envelope break down, the timing of which coincides with the end of phragmoplast expansion (Figure 3A). At the same time, SNAP-TM-mCherry disappeared from the plasma membrane adjacent to the cell plate (Figure 8C, white arrowheads), suggesting that SNAP-TM-mCherry accumulates at the cell plate by diffusing into the cell plate membrane from the parental plasma membrane once membrane fusion has occurred, rather than by delivery to the cell plate via exocytosis. By measuring the intensity of SNAP-TM-mCherry at the plasma membrane adjacent to the cell plate and in the middle of the cell plate, we quantified the kinetics of SNAP-TM-mCherry as it disappeared from the plasma membrane and appeared in the cell plate. Within 20 minutes after nuclear envelope break down, SNAP-TM-mCherry fluorescence decreased at the plasma membraned accompanied by an increase in the cell plate SNAP-TM-mCherry fluorescence. Interestingly *Δsabre* cells exhibited the same SNAP-TM-mCherry kinetics as wild type, suggesting that diffusion from the parental plasma membrane was not impaired and thus membrane fusion is unaffected in *Δsabre*.

To determine why cells that accumulated brown material often lysed, we used aniline blue staining to image accessible callose in cells with brown material. We found that compared to normal cell plates, which exhibited donut-shaped callose enrichments (Figure 8E), cells with brown material could be grouped into three categories: 27% stained weakly with aniline blue, 20% had a partial aniline blue ring, and 53% had large chunks of aniline blue staining near the brown material (Figure 8F; N=15 cells). Time-lapse imaging in a *Δsabre* cell that accumulated dark material and abnormal chunks of callose revealed accumulation of callose near the stalled microtubules that subsequently disappeared and reaccumulated on the other side of the cell (Figure 8G – video 14), suggesting that *Δsabre* cells exhibit defects in callose secretion and remodeling of callose during cell plate maturation.

## Discussion

The *SABRE* gene was identified nearly three decades ago. In the intervening time, SABRE has been found to influence cell growth and polarity in plants. Despite the rich phenotypic analyses of mutants in seed plants, uncovering the molecular basis of *SABRE* function has remained elusive. Here, we used a combination of genetics and live-cell imaging to study *SABRE* in the model bryophyte *P. patens*. Similar to *sabre* phenotypes in roots and shoots in Arabidopsis, *Δsabre* plants in moss were stunted, as a result of defects in cell expansion. Both polarized-growing protonemata and diffusely expanding cells in the phyllids of gametophores were smaller in *Δsabre* plants. Similar to defects in pollen tubes from mutants of *KIP*, a second *SABRE* gene in Arabidopsis (Xu and Dooner, 2006), we observed twisty protonemata that also had periods of normal growth interspersed with pauses. However, unlike pollen tubes or root hairs, moss protonemata also undergo cell division, which was dramatically affected in *Δsabre* plants. We discovered that 56% of seven-day-old plants regenerated from protoplasts exhibited at least one event where material that appears brown by bright field microscopy was deposited at the site of cell division (Figure 3). In extreme cases, defects during cell division led to failure characterized by loss of cell integrity and subsequent cell death. Interestingly, failures in cell division were readily detected in protonemata, but not in gametophores. In Arabidopsis, *sabre* mutant roots have defects in cell expansion and in placement of the cell division plane ultimately leading to dramatic defects in root morphology (Pietra et al., 2013). In contrast, shoot tissues were stunted but were morphologically normal (Pietra et al., 2013). Given that protonemata, which have more of a rooting function, exhibited stronger phenotypes than gametophores, the shoots of mosses, these results suggest that SABRE activity may play a more important role in rooting tissues than in shoots in land plants.

Current models of cell plate formation posit that Golgi- and trans-Golgi-derived vesicles accumulate at the phragmoplast midzone where they fuse to each other to form a tubular network whose lumen accumulates callose (Smertenko et al., 2017). As the phragmoplast expands, more vesicles are added to the edge and the tubular network enlarges. During expansion, clathrin-mediated endocytosis remodels the membrane transforming the tubular network into a fenestrated sheet, which coincides with the peak of callose accumulation (Samuels et al., 1995). Ultimately this sheet fuses with the parental cell membrane (Boruc and Van Damme, 2015; de Keijzer et al., 2017; Smertenko, 2018; Smertenko et al., 2017). Once fusion occurs the lumen of the cell plate becomes continuous with the apoplast, the extracellular environment of the plant, and the cell plate matures into a primary wall accompanied by changes in carbohydrate composition (Drakakaki, 2015). The exact mechanisms that dictate carbohydrate maturation are unclear but throughout cell plate formation numerous components of vesicle trafficking and endocytosis including have been shown to influence cell plate formation with distinct spatiotemporal contributions (Chow et al., 2008; Lauber et al., 1997; Rybak et al., 2014; Smertenko et al., 2017; Steiner et al., 2016). Microtubules and actin in the phragmoplast are hypothesized to direct vesicle trafficking as well as influencing cell plate positioning and structural stabilization of the nascent cell plate.

To determine how SABRE contributes to cell plate formation, we imaged cell division in *Δsabre* plants. While we observed normal actin and microtubule behaviors and dynamics during the initial stages of cytokinesis, we found that both actin and microtubules remained associated with the fully expanded cell plate for an extended period of time. During this stage of cytokinesis, the nascent cell plate matures into the cell wall. By following dividing cells, we discovered that during cell plate maturation aniline blue, a dye that binds to callose, takes significantly longer to accumulate in *Δsabre* cell plates as compared to wild type cell plates. Furthermore, membrane remodeling does not seem to be affected, since the behaviors of FM4-64 and a plasma membrane protein, SNAP-TM-mCherry, are similar in wild type and *Δsabre* cells. However, *Δsabre* cells that accumulated brown material exhibited a range of different callose staining behavior, from little staining to accumulations of large aggregates of callose. Time lapse imaging revealed that the large callose aggregates remodeled over time, suggesting that cells lacking SABRE exhibit unregulated callose deposition and remodeling. In seed plants, callose begins to accumulate during the ring phragmoplast stage (Park et al., 2014; Samuels et al., 1995), coincident with the timing of SABRE and ER recruitment along the cell plate (Figure 5). EM studies have shown that the ER accumulates parallel to either side of the developing cell plate once the fenestrated sheet has fully expanded (Seguí-Simarro et al., 2004). However, whether ER in this region simply repopulates a new cortical ER domain or if it plays a role in cell plate maturation has been unclear. Here, we provide striking evidence that the ER domains decorated with SABRE at the nascent cell plate play a critical role during cell plate maturation. Without SABRE, the ER still accumulates, but in a fully expanded phragmoplast, the ER parallel to the cell plate invariably buckles in the middle of the cell. Delayed callose deposition results in retention of both actin and microtubules at the expanded phragmoplast, perhaps to lend structural support to the nascent cell plate lacking callose. *Δsabre* cells that had unregulated callose deposition often accumulated brown material and many lost their integrity resulting in cell lysis and death, suggesting that the SABRE decorated ER domains contribute to properly regulating callose deposition. Whether this regulation is direct or via secretion of callose synthase proteins to the nascent cell plate remains to be determined.

Beyond cell division SABRE influences the directionality and persistence of polarized growth and nuclear migration. We discovered that SABRE forms dynamic puncta at the cell cortex and in the cytoplasm. With increased signal-to-noise, VAEM revealed that at the cortex, SABRE puncta moved along ER tubules. In Arabidopsis, a functional fluorescent fusion of SABRE, formed puncta throughout in the cytoplasm. Interestingly in this study, the authors were unable to determine co-localization to any specific organelle (Pietra et al., 2013). However, since the ER is such a complex network found throughout almost the entire cytoplasm, it might be difficult to identify co-localization to just a specific ER domain without the benefit of increased sensitivity afforded by VAEM imaging.

Given the striking co-localization of SABRE and fractions of the ER at the cell cortex, we investigated ER morphology in *Δsabre* in interphase. We discovered that the ER forms abnormal aggregates at the cell cortex and in the cytoplasm. Since the cytoskeleton is mostly unaffected in *Δsabre*, we hypothesize that nuclear migration and polarized growth defects stem from SABRE’s influence on the ER. Nuclear migration in the apical protonemal cell exhibits a distinct pattern including both apically and basally directed movements (Figure 6). While the physiological significance of these movements is unclear, studies in *P. patens* have identified microtubule motor proteins that mediate these nuclear migration events. Specifically, kinesin-14 drives basal movements (Yamada and Goshima, 2018), while kinesin-13 drives apical nuclear movement during prophase (Leong et al., 2020). Mutations in these motor proteins resulted in exaggerated movements in the opposite direction rather than a stationary nucleus, indicating that nuclear movement results from a balance of forces. In *Δsabre* we observed exaggerated basally directed nuclear movement during and after cell division and additionally, we found that interphase nuclei in the apical cell oscillated backwards as they moved apically towards the cell tip. Since *SABRE* influences ER morphology, we surmise that defects in nuclear migration stem from alterations to the nuclear envelope, which is contiguous with the ER. The exaggerated basal nuclear migration in *Δsabre* could be either the enhancement of basal moving forces or inhibition of the apical moving force, possibly generated by the endoplasmic reticulum or ER localized proteins.

How the ER might influence polarized growth persistence via *SABRE* is an interesting question. The ER accumulates just below the cell tip where both actin and microtubules drive and steer polarized growth, respectively in protonemata. However, in *Δsabre*, both the actin and microtubule cytoskeletons were not significantly altered, suggesting that SABRE’s impact on cell expansion is independent of the cytoskeleton. Of note, a recent study demonstrated that protonemata lacking COPII function, which mediates ER to Golgi transport, exhibited aggregated ER and polarized growth defects (Chang et al., 2020), suggesting that SABRE’s influence on the ER might alter ER secretory function. In contrast to defects in COPII function, which generally reduces secretion, *Δsabre* defects appear to specifically affect a subset of secretory cargo, given that delivery of the plasma membrane protein SNAP-TM-mCherry was unaffected in *Δsabre* but affected in COPII mutants (Chang et al., 2020). In a surprising connection, a study in *Drosophila* discovered hobbit, a protein that the authors report is conserved broadly across eukaryotes (Neuman and Bashirullah, 2018). *SABRE* is the putative plant hobbit homolog albeit with significant sequence divergence. Even with this divergence and the vast evolutionary distance between flies and plants, hobbit localizes to the ER when overexpressed in *Drosophila* and hobbit mutants are stunted similar to *sabre* null mutants in both Arabidopsis and *P. patens*. In *Drosophila*, mutants in hobbit accumulated proteins required for membrane fusion in endosomal compartments and were defective specifically in insulin secretion, manifesting in stunted growth. If hobbit and SABRE function are conserved, then in plants SABRE may regulate a subset of secretory cargos critical for cell expansion and division.

Alternatively, SABRE might influence the composition of regions of the ER membrane. Altered distribution or activity of ER resident membrane proteins, such as ethylene receptors (Ji and Guo, 2013; Yang et al., 2015) could impact growth and development. Previous studies in Arabidopsis provide a link between ethylene, a gaseous phytohormone involved in a variety of developmental processes and stress responses (Binder, 2020; Binder and Eric Schaller, 2017), and SABRE since inhibition of ethylene biogenesis partially rescued the *sabre* mutant in Arabidopsis (Aeschbacher et al., 1995; Yu et al., 2012). To distinguish between altered ethylene responses versus secretory defects, comparative RNA-seq and proteomic studies in *Δsabre* versus wild type could provide future research directions to narrow down SABRE’s influence on ER function. Another intriguing possibility is based on SABRE’s impact on callose deposition. Perhaps during cytokinesis, SABRE is recruited to the cell plate membrane via ER-plasma membrane contact sites and, there SABRE regulates callose synthase activity ensuring uniform deposition of callose.

Using a combination of live cell microscopy and genetics, we discovered that SABRE regulates ER morphology, which is contrary to previous work that had indicated *SABRE* influences microtubules (Pietra et al., 2013). Our results have revealed that the ER doesn’t simply repopulate at the daughter plasma membranes during cell division, but rather is critical for cell plate maturation and is involved in regulating callose deposition. Furthermore, the ER via SABRE, ultimately impacts cell expansion and nuclear migration. Future studies investigating interactions between *SABRE* and ER localized proteins involved in protein trafficking, ethylene sensing, and cell wall synthesis could distinguish whether the ER influence on cell division, cell expansion and nuclear migration result from defective secretion or altered ER membrane composition and function.

## Materials and Methods

### Plasmid construction

All genomic modifications in this study were performed using CRISPR-Cas9 mediated homology directed repair (HDR). In brief, two plasmids were generated: a CRISPR plasmid that contains the protospacer(s) and Cas9 ultimately generating the double-stranded break(s) at the designated genomic site(s), and a homology plasmid that provides the template in addition to the sequence being inserted (knockout cassette, fluorescent protein sequence, or promoter) for DNA repair. The two plasmids were co-transformed into moss protoplasts and transformants were regenerated from single protoplasts. All plasmids were constructed using the methods and modular vectors described in (Mallett et al., 2019). Primers used to generate these plasmids along with the corresponding plasmid products and primers used for subsequent genotyping are listed in Supplementary Table 1. Plasmids were transformed into a variety of moss lines stably expressing fluorescently labeled markers. These lines and the corresponding new lines generated in this study are listed in Supplementary Table 2. Note that for generating SABRE-3mNG tag, we initially transformed pMH-SAB-C and pGEM-SAB-3mNG into moss lines expressing lifeact-mRuby and mCherry-tubulin. To introduce ER labeling, we used CRISPR mediated HDR, to swap lifeact-mRuby into mCherry-KDEL (Supplementary Table 2). All other fluorescent labeled lines are as described before: GFP-tubulin (Wu and Bezanilla, 2018), Lifeact-GFP (van Gisbergen et al., 2012), SP-GFP-KDEL (Chang et al., 2020), NLS-GFP-GUS/SNAP-TM-mCherry (van Gisbergen et al., 2018, p. 10), mCherry-tubulin (Burkart et al., 2015), Lifeact-mRuby (van Gisbergen et al., 2020).

To generate the three tandem mNeonGreen (3mNG) tag, we amplified mNG codon optimized for budding yeast with DC625 and DC626 (Supplemental Table 1) incorporating a BamHI site just upstream of the ATG. This PCR product was introduced into pDONR221-P4rP3r using BP reaction (Invitrogen) generating pENTR-R4R3-mNG. A second mNG codon optimized for *P. patens* was amplified without the stop codon and incorporating BamHI upstream and BglII downstream. This product was ligated into pENTR-mNG in-frame upstream using the BamHI site to create pENTR-R4R3-2XmNG. The third mNG codon optimized for budding yeast was similarly amplified and ligated into pENTR-R4R3-2XmNG. The resulting pENTR R4R3 Cterm BamHI 3xSc_Pp_Sc_mNeon plasmid (Supplementary Table 1) was used to create the homology plasmid pGEM-SAB-3mNG according to (Mallett et al., 2019). Similarly, to knock-in the stronger constitutive maize ubiquitin promoter, pENTR-R4R3-Ubiquitin-pro was generated by inserting the ubiquitin promoter into pDONR221P4rP3r with BP reaction (Supplementary Table 1), and the resulting entry clone was used to make the homology plasmid for moss transformation.

### Plant culture and transformation

Moss tissue was cultured on PpNH4 medium (1.03 mM MgSO4, 1.86 mM KH2PO4, 3.3 mM Ca(NO3)2, 2.72 mM (NH4)2-tartrate, 45 μM FeSO4, 9.93 μM H3BO3, 220 nM CuSO4, 1.966 μM MnCl2, 231 nM CoCl2, 191 nM ZnSO4, 169 nM KI, and 103 nM Na2MoO4) supplied with 0.7% agar, plated on petri dishes. After propagation by blending, 5 to 7-day old tissue was protoplasted and then transformed as previously described (Liu and Vidali, 2011). For all the HDR transformations, 7.5 μg of the CRISPR/protospacer plasmid and 7.5 μg of the homology plasmid were co-transformed into 150 μL protoplasts at a concentration of 2,000,000 protoplasts/mL. Transformed protoplasts were resuspended in liquid plating medium (PpNH4 plus 8.5% mannitol and 10mM CaCl2), plated and regenerated on PRM-B media (PpNH4 plus 6% mannitol and 10mM CaCl2) with 0.8% agar. A layer of cellophane was placed on top of the PRM-B plates and protoplasts were plated on top of the cellophane. After 4 days on PRM-B, the cellophane was transferred to PpNH4 supplied with antibiotic for selection. 15 μg/mL Hygromycin was used for selection of transformed protoplasts. Plants were grown on selection for a week before moving to PpNH4 media for subsequent culturing and genotyping.

Growth assays were used to quantify protonemal area. Tissue regenerated from protoplasts was used to synchronize plant growth. Protoplasts were isolated, plated and regenerated as described above. After 4 days on PRM-B, they were transferred to PpNH4 and allowed to grow for another 3 days. Seven days after protoplasting, plants were imaged with a Nikon SMZ25 stereomicroscope equipped with a color camera (Nikon digital sight DS-Fi2). Plants were transferred from the plate to a slide and stained with 0.1 mg/mL calcofluor. Calcofluor fluorescence was imaged with a violet filter cube (excitation 420/25, dichroic 455, emission 460 longpass). Subapical cell length was measured manually using these images. Quantification of plant area was carried out using methods modified from (Vidali et al., 2007). In brief, colored images were converted to a single red color image. Single plants were selected and highlighted by cropping and thresholding above a certain intensity value. Plant area was calculated based on the thresholded images. For each experiment, plant area was normalized to the average area of control plants.

### Brightfield microscopy

Brightfield time-lapse microscopy was performed using a Nikon Ti microscope equipped with a 0.8NA 20× objective. Plants were cultured in continuous light in PDMS microfluidic devices with liquid Hoagland’s medium as previously described (Bascom et al., 2016). For imaging protonemal tissue, ground tissue was loaded into microfluidic devices and allowed to grow at least 2-3 days before imaging. Gametophores emerged from protonemata naturally 2-3 weeks after loading ground tissue. Multiple XY positions were acquired, and a either a single focal plane or a z-stack was acquired for each position. Between each time point, white light remained on to provide light for plant growth. Mono-color bright field images were acquired with Nikon DS-Qi2 camera. Colored bright field images were acquired with Nikon DS-Vi1 camera. Extended-depth of focus images were created for the Z-stacks with NIS-element (Nikon). Protonemata growth rate was measured by manually tracing the growing tip during active growing period in NIS-element software.

### Fluorescence microscopy

For short-term imaging, moss protonemal tissue was mounted on an agar pad on a slide, submerged in Hoagland’s medium and sealed with a coverslip. For protonemata staining, FM4-64 (15 μM), aniline blue (20 μg/mL) and Fast Scarlet (50 μg/mL) were dissolved in Hoagland’s medium used for tissue mounting. For imaging tissue from regenerated from protoplasts, regenerating protoplasts were removed from the cellophane, loaded into microfluidic devices and immediately imaged. To synchronize cell divisions, ground tissue was loaded into microfluidic devices and allowed at least 4 days to grow. The microfluidic device was placed in far red light for 3-4 days before exposure to white light and imaging. Confocal imaging was performed using a Nikon A1R laser scanning confocal with a 1.3 NA 40× or 1.49 NA 60× oil immersion objective (Nikon). Laser illumination at 405nm was used for exciting aniline blue dye, 488 nm was used for exciting mNeonGreen, GFP and chlorophyll autofluorescence; 561 nm for mRuby2, mCherry, FM4-64, Fast Scarlet and propidium iodide. Emission filters were 525/50 nm for mNeonGreen/GFP and aniline blue; 595/50 nm for mRuby2, mCherry, FM4-64, Fast Scarlet and propidium iodide. For chlorophyll autofluorescence emission light passed through a long-pass filter allowing wavelengths larger than 640 nm to pass. Image acquisition was controlled by Nikon NIS-Element software (Nikon). In between each time point, transmitted white light was on providing light for plant growth. 3D reconstruction was done using 3D volume viewer with maximum projection rendering method in NIS-Element, contrast for slices at different Z positions were adjusted individually to compensate for loss of signal in tissue further away from the objective using 3D lookup table function. Deconvolution was carried out with NIS-Elements (Nikon) with the 2D deconvolution default settings.

To quantify cell size in gametophores, mature phyllids were removed from gametophore and mounted in a droplet of staining solution (15μg/mL propidium iodide dissolved in liquid Hoagland’s medium) between a slide and coverslip. Confocal images were captured for quantification. In Fiji (Schindelin et al., 2012), fluorescent images were processed using enhance contrast, subtract background, smooth and median filter. The processed images were then converted to a binary mask and put through binary process – Close>Dilate>Close>Skeletonize>Dilate, to outline the edges of the cells. Images were then inverted to highlight the cell area, and subsequently quantified using the analyze particle function. After quantification, incorrect cells (fused or broken) were manually removed.

Variable-angle epifluorescence microscopy (VAEM) microscopy was performed using a Nikon Ti-E inverted microscope equipped with a TI-TIRF-PAU illuminator, using a Nikon 1.49 NA 100x oil immersion TIRF objective. 6 to 8-day old plants regenerated from protoplasts were mounted between a coverslip and an agar pad on top of a slide, prepared right before imaging. GFP and mNeonGreen were illuminated with a 488nm laser, while mRuby2 and mCherry were excited with a 561nm laser, the emission passed through a 525/50 filter for GFP/mNeonGreen and 610/75 for mRuby2/mCherry. Images were simultaneously captured with two Andor 897 EMCCD cameras. Image acquisition was controlled by Nikon NIS-Elements software. All data was processed with enhanced contrast (0.1% pixel saturation), subtract background and smoothing in Fiji using default settings. Pearson’s correlation coefficients were calculated in NIS-Elements. Cortical actin and microtubule dynamics were quantified by measuring the decay of the correlation coefficient over time, as previously described (Vidali et al., 2010).

### Fluorescence recovery after photobleaching (FRAP)

Phragmoplast microtubule photobleaching experiments were conducted using a Nikon A1R laser scanning confocal microscope with 1.49 NA 60× oil immersion objective. Actively growing plant tissue was mounted on an agar pad between the slide and coverslip, then imaged immediately with a 488 nm laser to identify actively dividing cells at the phragmoplast expansion stage. A 3 μm X 3 μm square region of interest (ROI) was placed in the center of phragmoplast, ensuring the entire ROI was filled with phragmoplast microtubules. Photobleaching was carried out using a 405 nm laser at 10% power for 1 second, after 6 frames (5 seconds) of normal imaging. Imaging continued after photobleaching for two minutes to capture fluorescence recovery. The average intensity in the ROI was measured for each frame, then normalized to the average of the value from the first 6 frames.

### Nuclear migration trajectory analysis

A moss line with the nuclear marker NLS-GFP-GUS and the plasma membrane marker SNAP-TM-mCherry (van Gisbergen et al., 2018) was imaged with time-lapse confocal microscopy. A z-stack was taken every 5 minutes for the apical cells of several filaments. A segmented line was drawn manually along the axis of growth to generate the kymograph. To make the trends on the kymograph easier to label (Figure 9C), we stretched the kymograph image by increasing the Y axis 2-fold. To measure the basal nuclear position, using the straight-line tool in Fiji we measured the distance between the middle of the cell plate and the basal edge of the nucleus when it was closest to the cell plate in the kymograph. The distance was divided by the cell length at the same time point, to generate the relative basal nuclear position. To analyze nuclear migration, we isolated movie fragments of nuclear apical migrating periods, tracked the nuclear GFP signal with the TrackMate plugin in Fiji (Tinevez et al., 2017), with a spot diameter of 10 μm. Displacement, distance and velocity were calculated. Nuclear migration displacement was defined as the straight-line distance between the initial and final positions of the nucleus. Total migration distance was calculated as the sum of displacement between each frame. Average instantaneous velocity was calculated between each frame.

## Acknowledgements

We thank Charles Barlowe lab (Dartmouth College) and Gohta Goshima lab (Nagoya University) for sharing mNeonGreen gene template with two different codon usages. We thank Ann Lavanway from Dartmouth College for helping with VAEM (TIRF) microscopy. The acquisition of the TIRF microscope at Dartmouth College was funded by an NIH S10, grant number 1S10OD018046-01. This work was supported by Dartmouth College, a grant from the National Science Foundation (MCB-1715785 to M.B.) and the John H. Copenhaver Jr. and William H. Thomas MD 1952 Award from the Dartmouth Molecular and Cellular Biology program (X.C.).

## Video Legends

**Video 1. Early gametophore development in wild type and *Δsabre*.** Each frame is an EDF image generated from a bright field Z-stack taken every 15 minutes. Scale bar, 10 μm. Video is playing at 10 frames per second (fps).

**Video 2. *SABRE* influences polarized growth in protonemata.** Bright field images were taken every 10 minutes. Scale bar, 5 μm. Video is playing at 10 fps.

**Video 3. *SABRE* influences diffused growth in gametaphores.** EDF images of bright field Z-stacks taken every hour. Scale bar, 20 μm. Video is playing at 5 fps.

**Video 4. Delays in disassembling phragmoplast microtubules in *Δsabre* mutant.** Images are maximum projections of confocal Z stacks of GFP-tubulin (green) and chlorophyll autofluorescence (magenta) in control and *Δsabre* cells acquired every 2 minutes. Video is playing at 5 fps. Scale bar, 10 μm.

**Video 5. Brown material deposition in *Δsabre* occurs during cell plate formation.** Each frame is a bright field EDF image taken every 10 minutes. Video is playing at 8 fps. Scale bar, 5 μm.

**Video 6. Representative VAEM time-lapse acquisitions showing localization of SABRE with microtubules, actin, or ER.** SAB-3mNG or OE-SAB-3mNG (green), and ER (mCherry-KDEL), microtubules (mCherry-tubulin) or actin (LA-mRuby) (magenta) are shown. Time-lapse acquired every 120 ms. Video is playing at 10 fps. Scale bar, 2 μm.

**Video 7. SABRE accumulation correlates with ER localization at the phragmoplast and both signals accumulated during later stages of cell division.** Images are single focal planes in the medial section of the cell. OE-SAB-3mNB (SAB), mCherry/GFP-tubulin (MT), ER-mCherry (ER). Frame interval is 1 minute. Video is playing at 5 fps. Scale bar, 3 μm.

**Video 8. *Δsabre* exhibits bright ER aggregates in the cytoplasm.** Images are single focal planes in the medial section and maximum projections of confocal Z-stacks of the cells showing SP-GFP-KDEL in control and *Δsabre*. Confocal Z-stacks acquired every 10 minutes. Video is playing at 4 fps. Scale bar, 5 μm.

**Video 9. ER buckles during cell plate formation in *Δsabre*.** Each frame is a single focal plane in the medial section of the cell, taken every 3 minutes. Magenta arrow indicates buckling. Video is playing at 5 fps. Scale bar, 5 μm.

**Video 10. Time lapse imaging showing exaggerated nuclear basal migration in relation to phragmoplast ER buckling during cell plate formation.** *Δsabre* cell 1 showed delayed apical nuclear movement in phase I and exaggerated basal movement in phase II. *Δsabre* cell 2 showed only exaggerated basal movement in phase II. Both cells exhibited ER buckling at the cell plate. Images are single confocal images taken every 10 minutes. Video is playing at 4 fps. Scale bar, 5 μm.

**Video 11. Nuclear movement during tip growth and cell division.** Blue traces represent the mother nuclei, red and green traces represent the daughter nuclei in the apical and sub-apical cells, respectively. Nuclei are shown in green or grey with colored tracks, and plasma membrane shown in magenta. The column on the far right depicts a *Δsabre* cell showing impaired forward nuclear movement before cell division and exaggerated basal nuclear movement after cell division. The nucleus moved basally all the way to the newly formed cell plate, appearing to smash into the cell plate, with an evident change in nuclear shape. The nucleus flattens out when in contact with the cell plate, observed at 510 to 525min. Video is a maximum projection of confocal Z-stacks taken every 5 min. Video is playing at 8 fps. Scale bar, 10 μm.

**Video 12. ER, actin and microtubule behavior during brown material deposition and cell division failure in *Δsabre*.** Single focal plane of ER (SP-GFP-KDEL), maximum projection of actin (Lifeact-GFP) and microtubules (GFP-tubulin) are shown. Time-lapse imaging was acquired every 10 minutes. Video is playing at 5 fps. Scale bar, 10 μm.

**Video 13. Aniline blue and FM4-64 staining at developing cell plate.** Green, FM4-64. Magenta, aniline blue. Images are medial sections from dividing cells. Scale bar, 5 μm. Video is playing at 8 fps.

**Video 14. Aniline blue staining during normal division and cytokinesis failure in *Δsabre*.**

Images are maximum projections of deconvolved confocal z-stacks taken every 10 minutes. Microtubule, green. Aniline blue, magenta. Scale bar, 10 μm. Video is playing at 5 fps.

**Supplementary Table 1.**
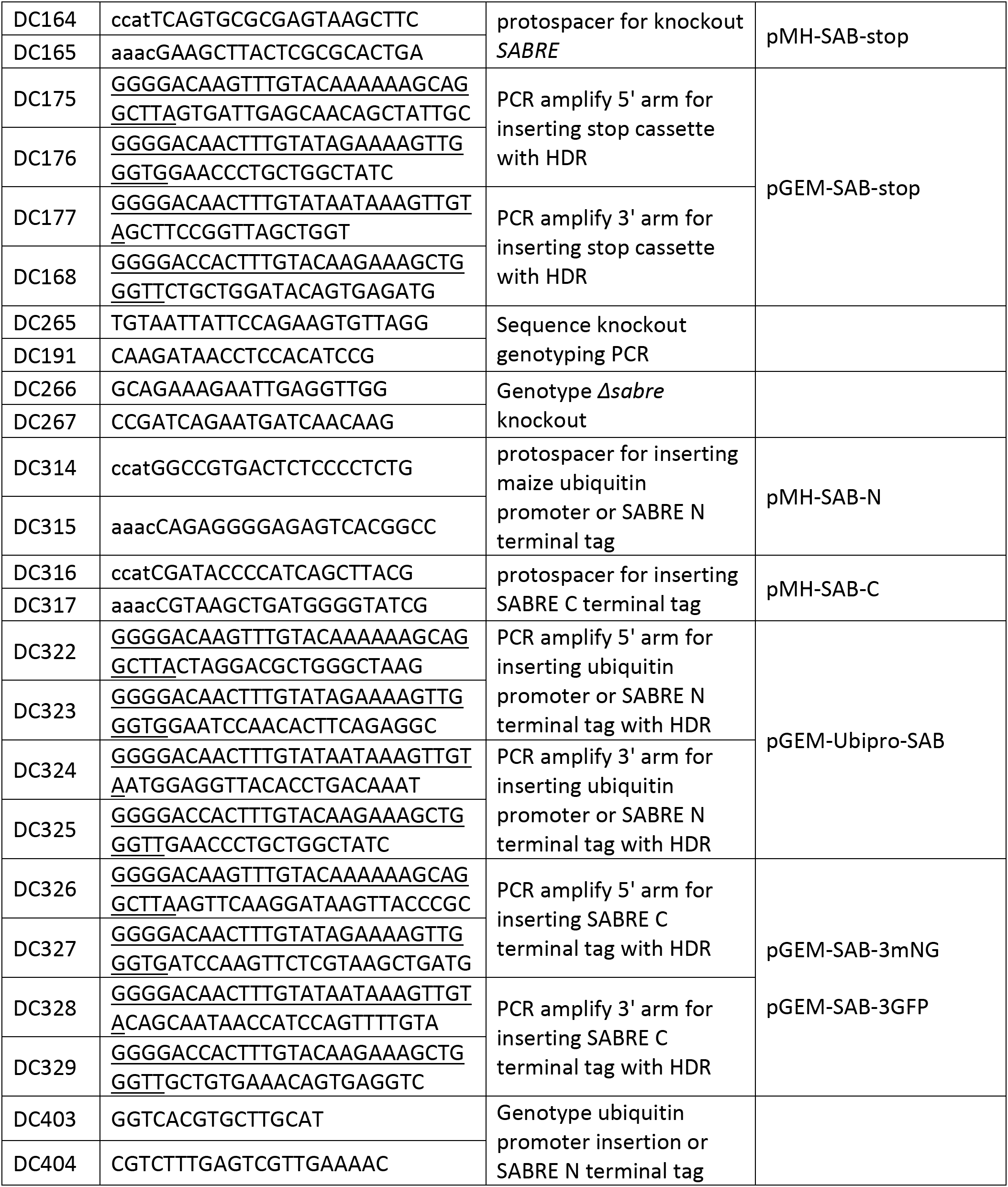
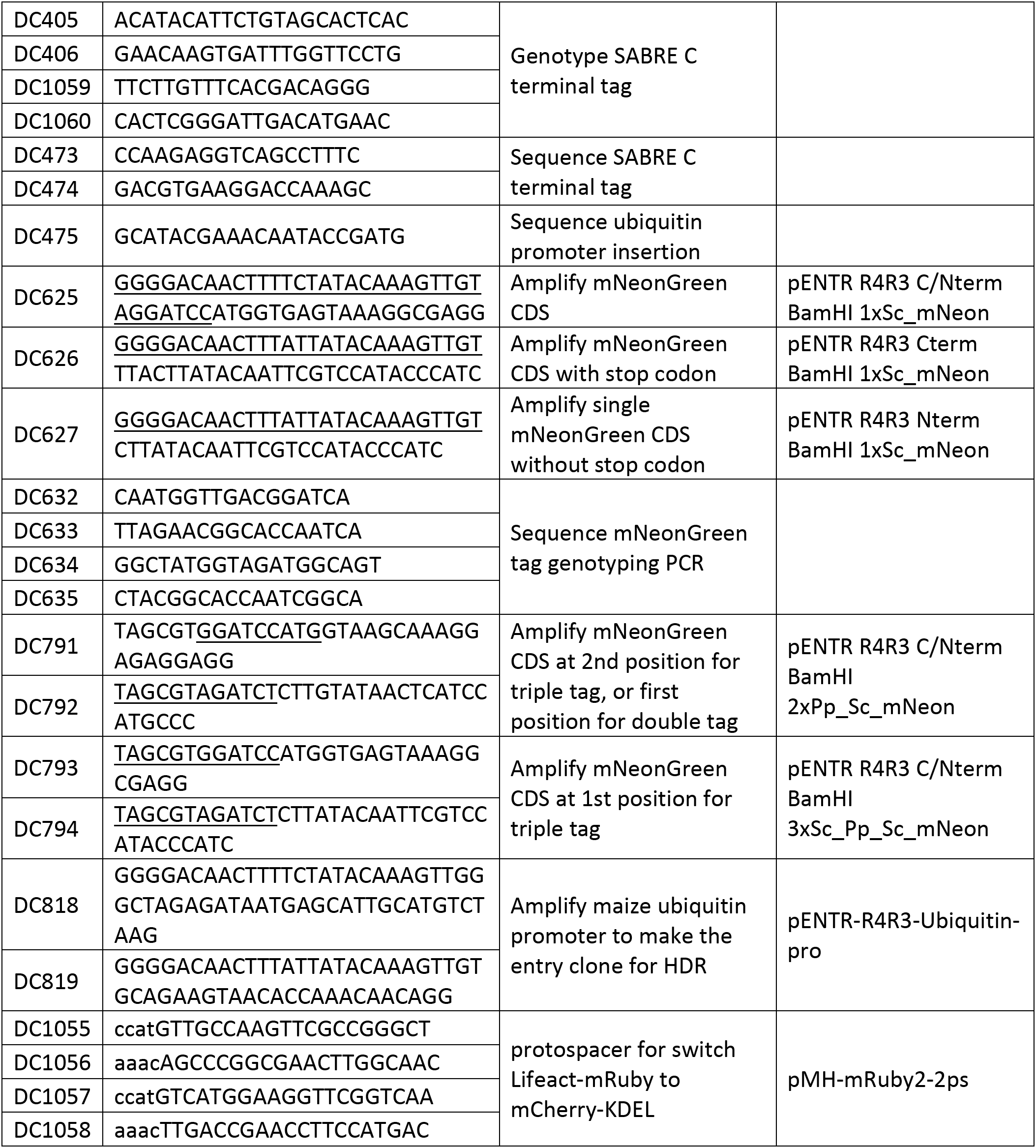
Primers used in this study and the plasmid constructs they are used to generate respectively

**Supplementary Table 2.** Plasmids used to transform moss and the lines generated from those transformations.

| **CRISPR plasmid** | **Homology plasmid** | **Background** | **Generated line** |
| --- | --- | --- | --- |
| pMH-SAB-stop | pGEM-SAB-stop | wild type | *Δsab*/wildtype |
| pMH-SAB-stop | pGEM-SAB-stop | GFP-tubulin | *Δsab*/GFP-tubulin |
| pMH-SAB-stop | pGEM-SAB-stop | Lifeact-GFP | *Δsab*/Lifeact-GFP |
| pMH-SAB-stop | pGEM-SAB-stop | SP-GFP-KDEL | *Δsab*/ER-GFP |
| pMH-SAB-stop | pGEM-SAB-stop | NLS-GFP-GUS/SNAP-TM-mCherry | *Δsab*/NLS-GFP/SNAP-mCh |
| pMH-SAB-C | pGEM-SAB-3GFP | wild type | SAB-3GFP |
| pMH-SAB-C | pGEM-SAB-3mNG | mCherry-tubulin | SAB-3mNG/mCh-tub |
| pMH-SAB-C | pGEM-SAB-3mNG | Lifeact-mRuby | SAB-3mNG/LAmR |
| pMH-SAB-N | pGEM-Ubipro-SAB | SAB-3mNG/mCh-tub | OE-SAB-3mNG/mCh-tub |
| pMH-SAB-N | pGEM-Ubipro-SAB | SAB-3mNG/LAmR | OE-SAB-3mNG/LAmR |
| pMH-mRuby2-2ps | pTZ-SP-mCherry-KDEL | SAB-3mNG/LAmR | SAB-3mNG/ERmCh |
| pMH-mRuby2-2ps | pTZ-SP-mCherry-KDEL | OE-SAB-3mNG/LAmR | OE-SAB-3mNG/mCh-tub |

**Figure 1 – figure supplement 1.**
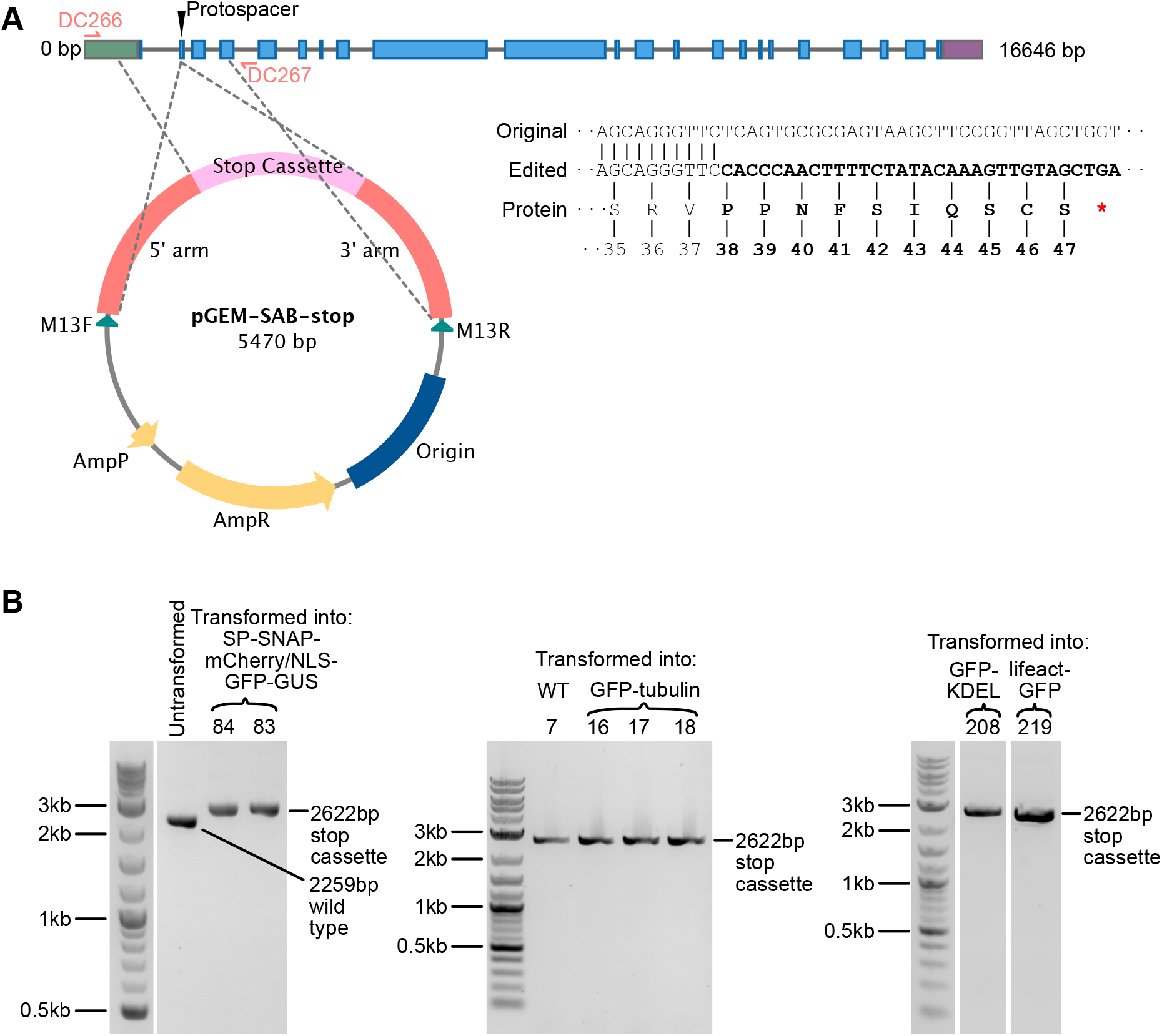
Generating SABRE null mutant with stop cassette insertion. (A) Gene model of *P. patens SABRE* and scheme for generating the null mutation. Protospacer indicates the region targeted for double stranded break and insertion of the stop cassette. Grey line, introns; green box, 5’UTR; purple box, 3’UTR; blue boxes, exons. Grey dashed lines extending from the plasmid map indicate the regions of homology. Original and edited DNA sequences near the insertion site are shown. Bold font indicates DNA sequences of inserted stop cassette and its translated protein that is different from the original sequence. Red asterisk indicates the premature stop codon. (B) Gel images showing PCR products amplified from mutants isolated in a variety of different backgrounds with the HDR stop cassette insertion amplified by primer pairs DC266, DC267. An untransformed control is shown in the first gel. All insertions were validating by sequencing.

**Figure 1 – figure supplement 2.**
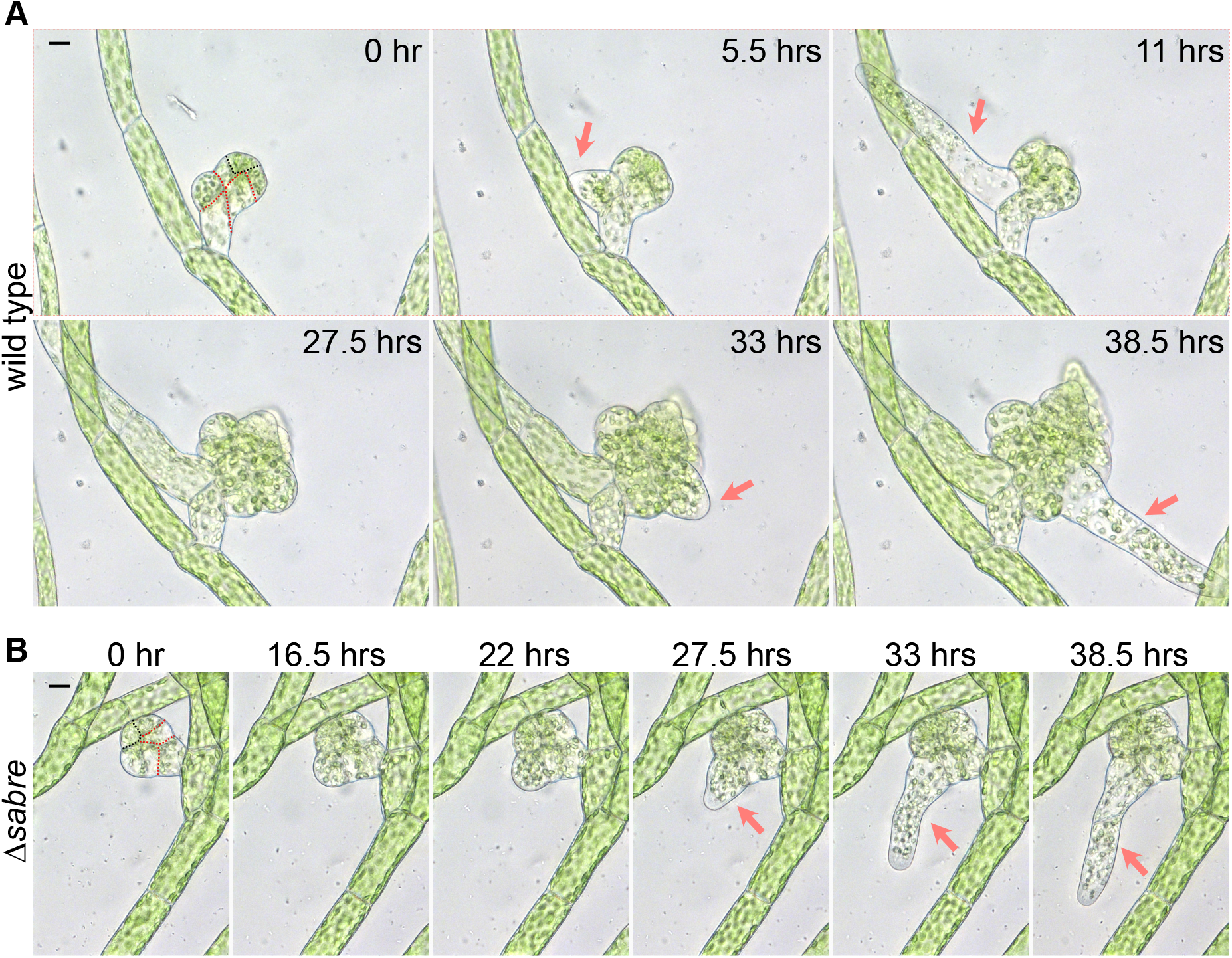
Early gametophore development. Gametophore initials (buds) from wild type (A) and *Δsabre* (B). Both have undergone stereotypic divisions (dashed red lines) to produce the apical pyramidal stem cell (dashed black line). In wild type (A), two rhizoids (arrows) emerged from the base of the developing gametophore and elongated by polarized growth. Scale bar, 10μm. A *Δsabre* (B) gametophore of similar age to the wild type shown in (A) growing for the same period of time. The *Δsabre* bud and rhizoid (arrows) grew significantly less than wild type. Scale bar, 10μm. Also see Video 1.

**Figure 2 – figure supplement 1.**
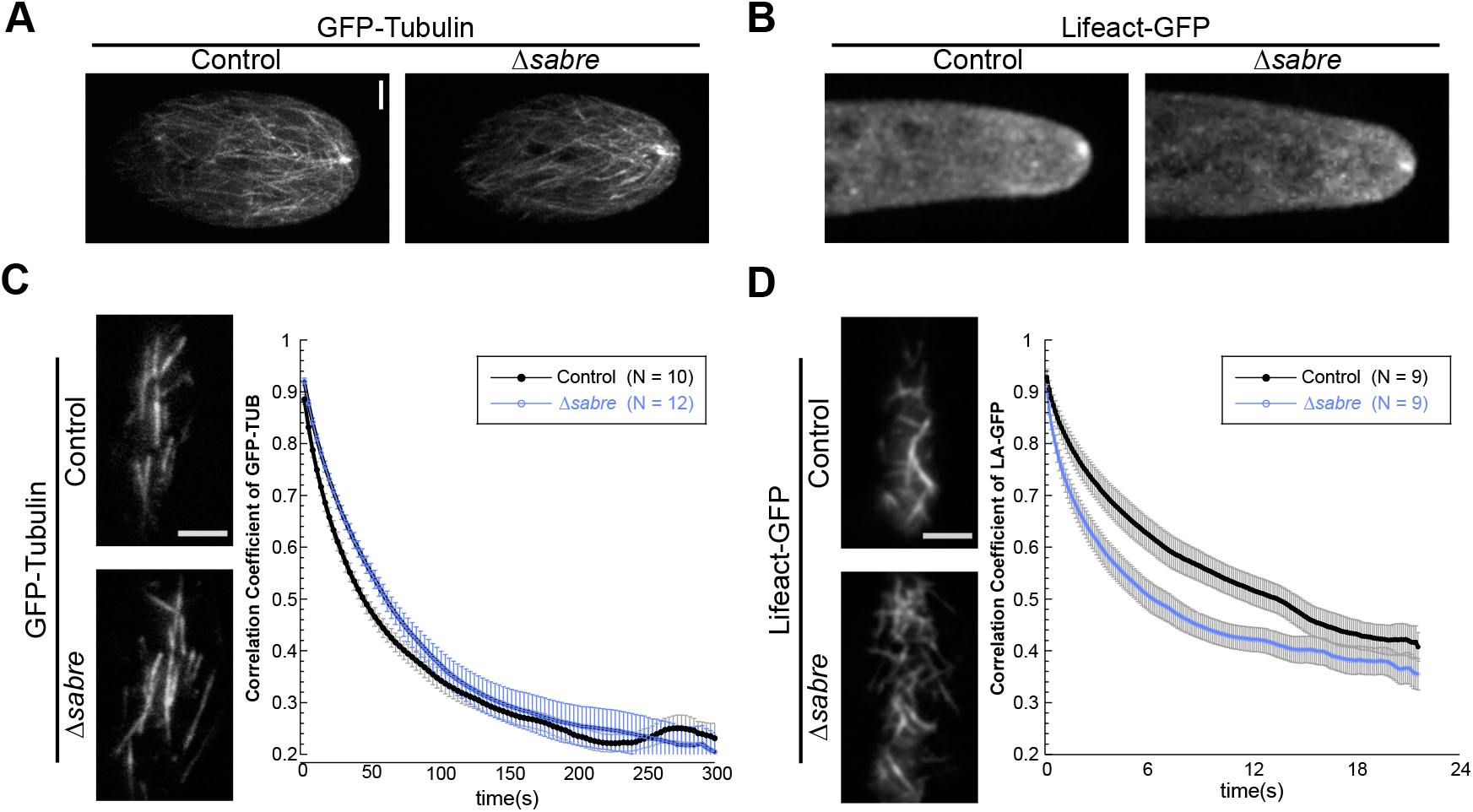
Organization and dynamics of actin and microtubule cytoskeletons in *Δsabre*. (A) Microtubule and (B) actin tip foci are not altered in *Δsabre*. Images are maximum projections of confocal Z-stacks. Scale bar, 3 μm. (C-D) Quantification of cortical microtubule (C) and actin (D) dynamics. Representative images are single frames of a VAEM time-lapse acquisition showing the over all cortical cytoskeleton organization. Graphs depict the correlation coefficient between images overall possible temporal spacings (time interval, X axis). Each data point is the average of several cells, Number of cells for each category indicated in graph legends. Scale bars, 3 μm. Error bar, standard error of the mean.

**Figure 3 – figure supplement 1.**
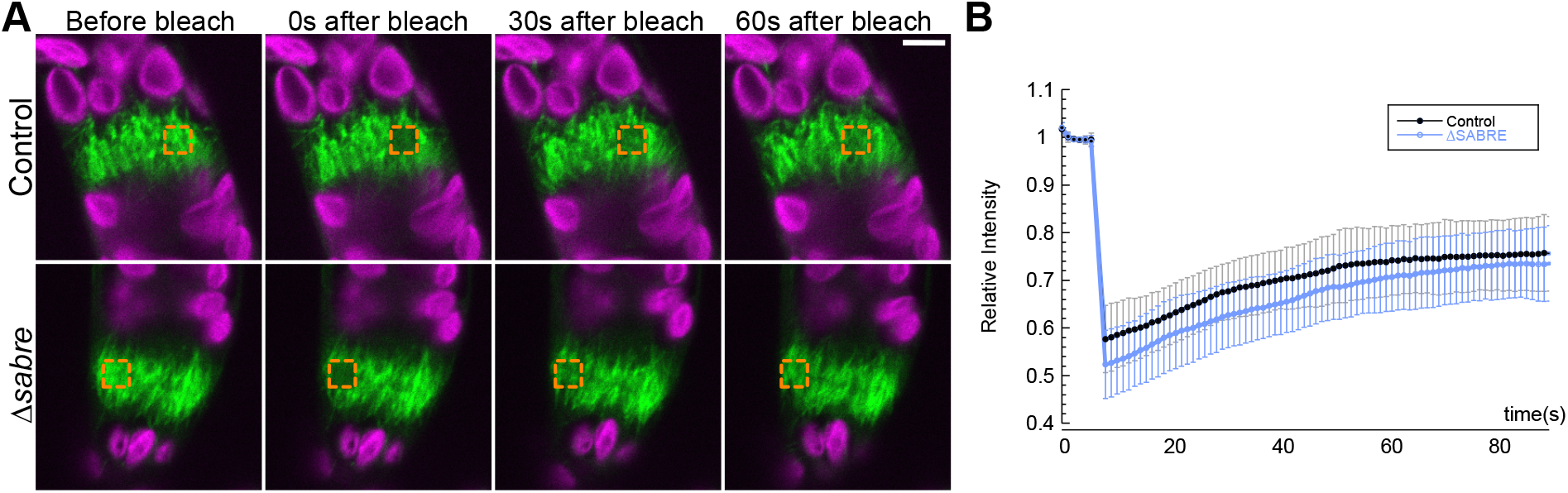
Phragmoplast microtubule dynamics are not altered in *Δsabre*. (A) Photobleaching of phragmoplast microtubules labeled with GFP-tubulin. Orange square indicates photobleached region. GFP-tubulin (green) and chlorophyll autofluorescence (magenta) are shown. Scale bare, 5 μm. (B) Relative intensity is plotted against time to show the recovery rate of fluorescence over the following 1.5 minutes after the bleaching event. Error bars represent standard deviation.

**Figure 4 – figure supplement 1.**
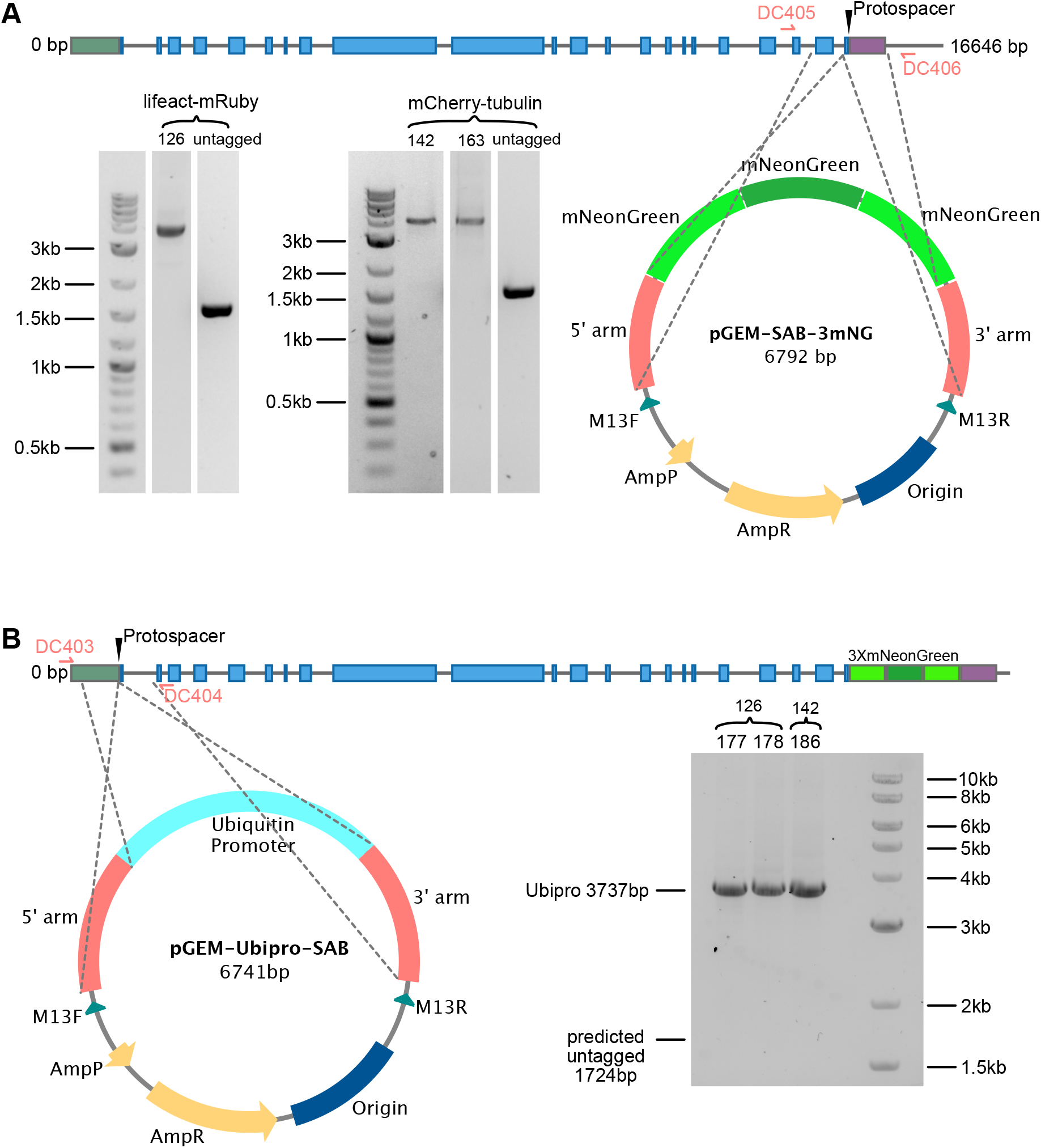
Model for generating SABRE-3mNG tag and overexpression with maize ubiquitin promoter. (A-B) Generating (A) 3mNG tag at C terminus and (B) inserting maize ubiquitin promoter at N terminus of SABRE. Protospacer indicates the region targeted for double stranded break. Grey lines, introns; green box, 5’UTR; purple box, 3’UTR; blue boxes, exons. Grey dashed lines extending from the plasmid maps indicate the regions of homology. Gel images shows PCR products amplified from transformed plants amplified by primer pairs DC405, DC406 for (A), and DC403, DC404 for (B). All insertions were validating by sequencing.

**Figure 4 – figure supplement 2.**
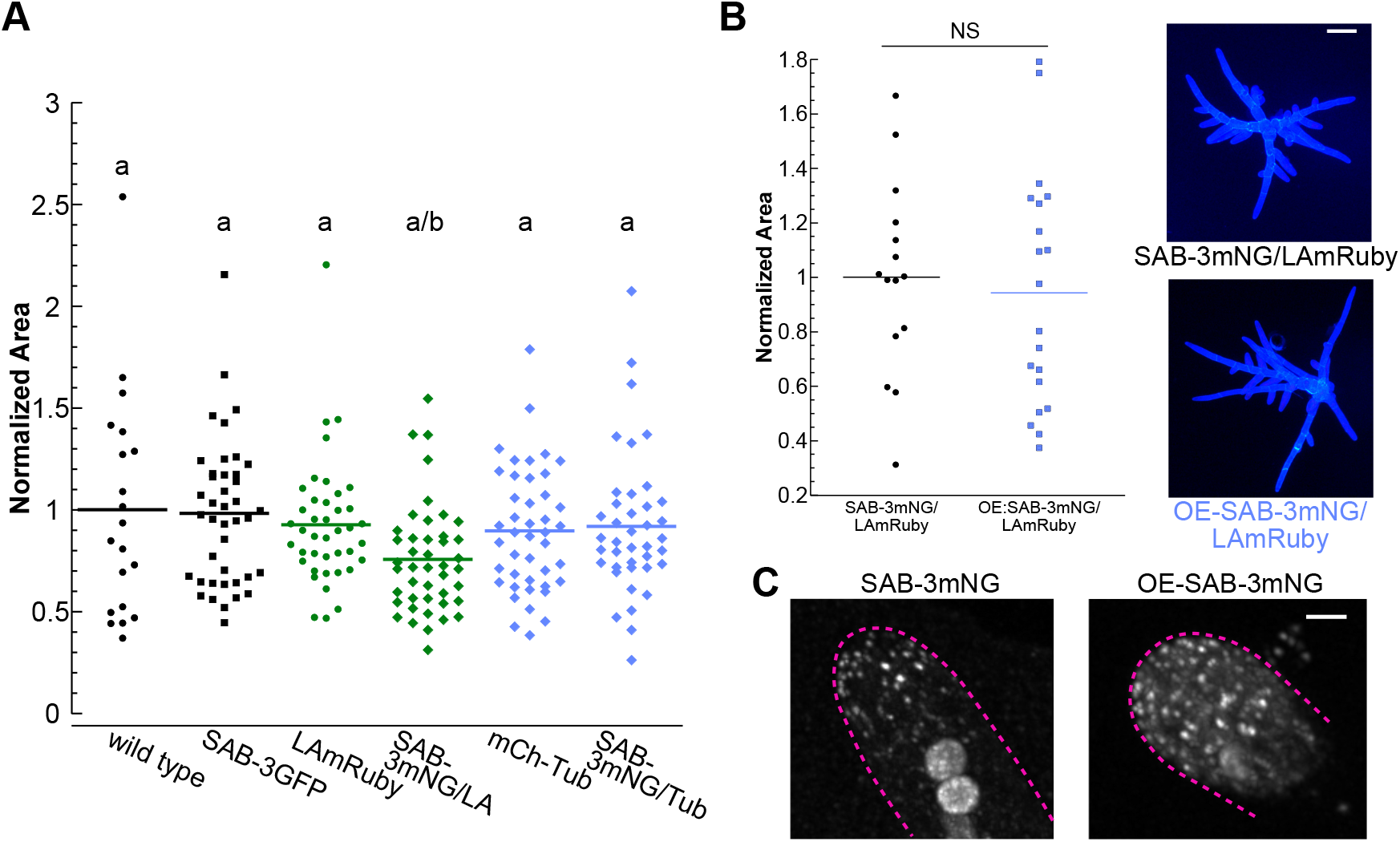
C-terminal tagging and over expression of SABRE does not influence protein function. (A) Quantification of plant area for the indicated genotypes of 7 day-old plants regenerated from protoplasts. Plant area was normalized to wild type. N=20, wild type; N=41, SAB-3GFP; N=41, LA-mRuby; N=44, SAB-3mNG/LA-mRuby; N=43, mCherry-Tub; N=40, SAB-3mNG/mCherry-Tub. Letters indicate groups with significantly different means as determined by ANOVA with a Tukey’s HSD all pair comparison post-hoc test (α=0.05). (B) Comparison of SAB-3mNG and OE-SAB-3mNG plant area. Plant area was normalized to SAB-3mNG. N=15, SAB-3mNG; N=20, OE-SAB-3mNG. A Student’s t-test for unpaired data with equal variance was performed. NS: not significant. Right panel, representative images of 7 day-old plants regenerated from protoplasts of the indicated genotype. Scale bar, 100 μm. (C) Example images comparing the density of SAB-3mNG and OE-SAB-3mNG. Magenta dashed lines label the cell outlines. Scale bar, 3 μm.

**Figure 5 – figure supplement 1.**
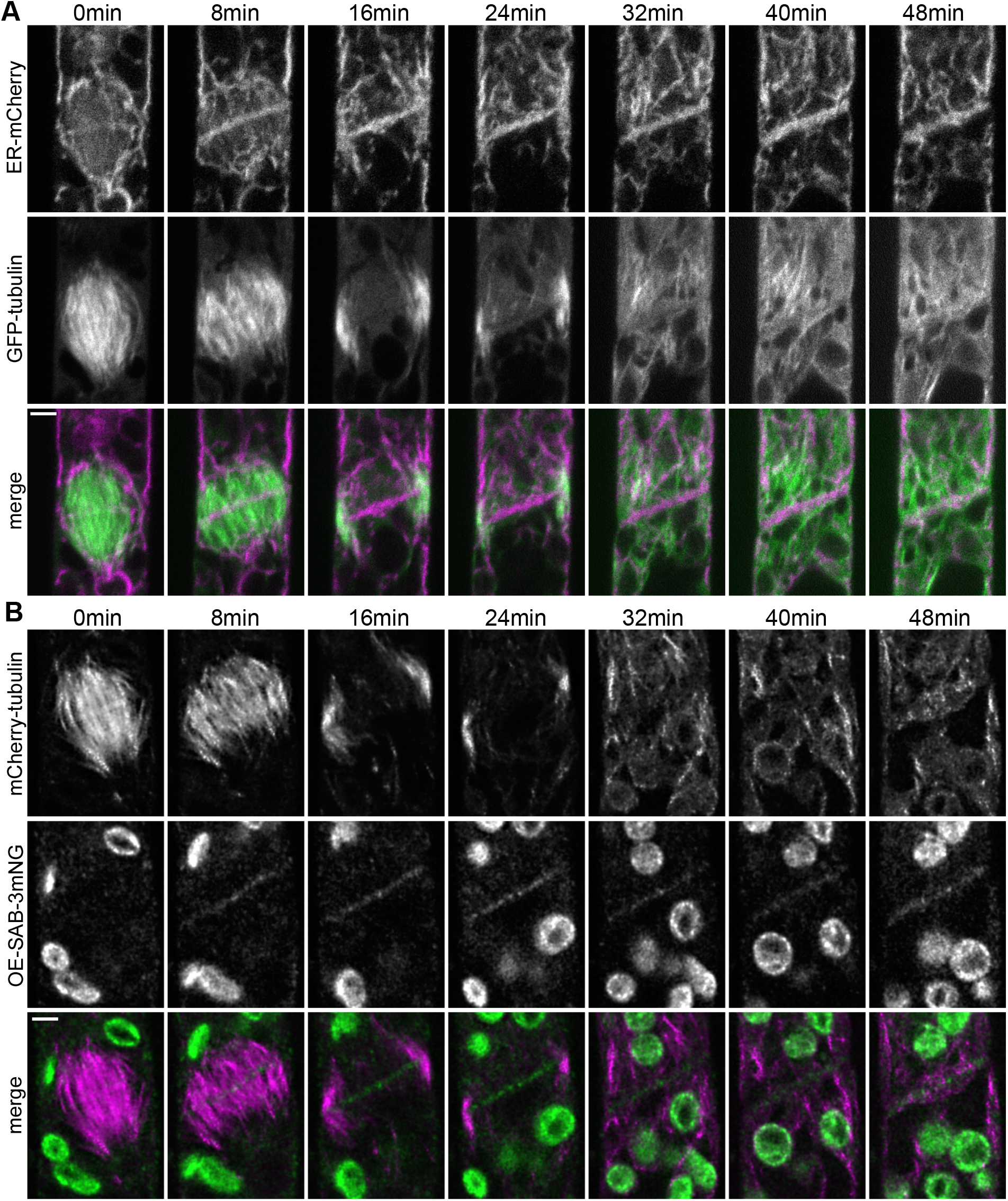
Timing of ER, microtubule and SABRE localization during cell plate formation. (A) Confocal images of GFP-tubulin (green in merge) and ER-mCherry (magenta in merge) during cell division in a wild type cell. Scale bar, 3 μm. Also see Video 7. (B) Deconvolved images of OE-SAB-3mNG (green in merge) and mCherry-tubulin (magenta in merge) during cell division. Scale bar, 3 μm. Also see Video 7.

**Figure 6 – figure supplement 1.**
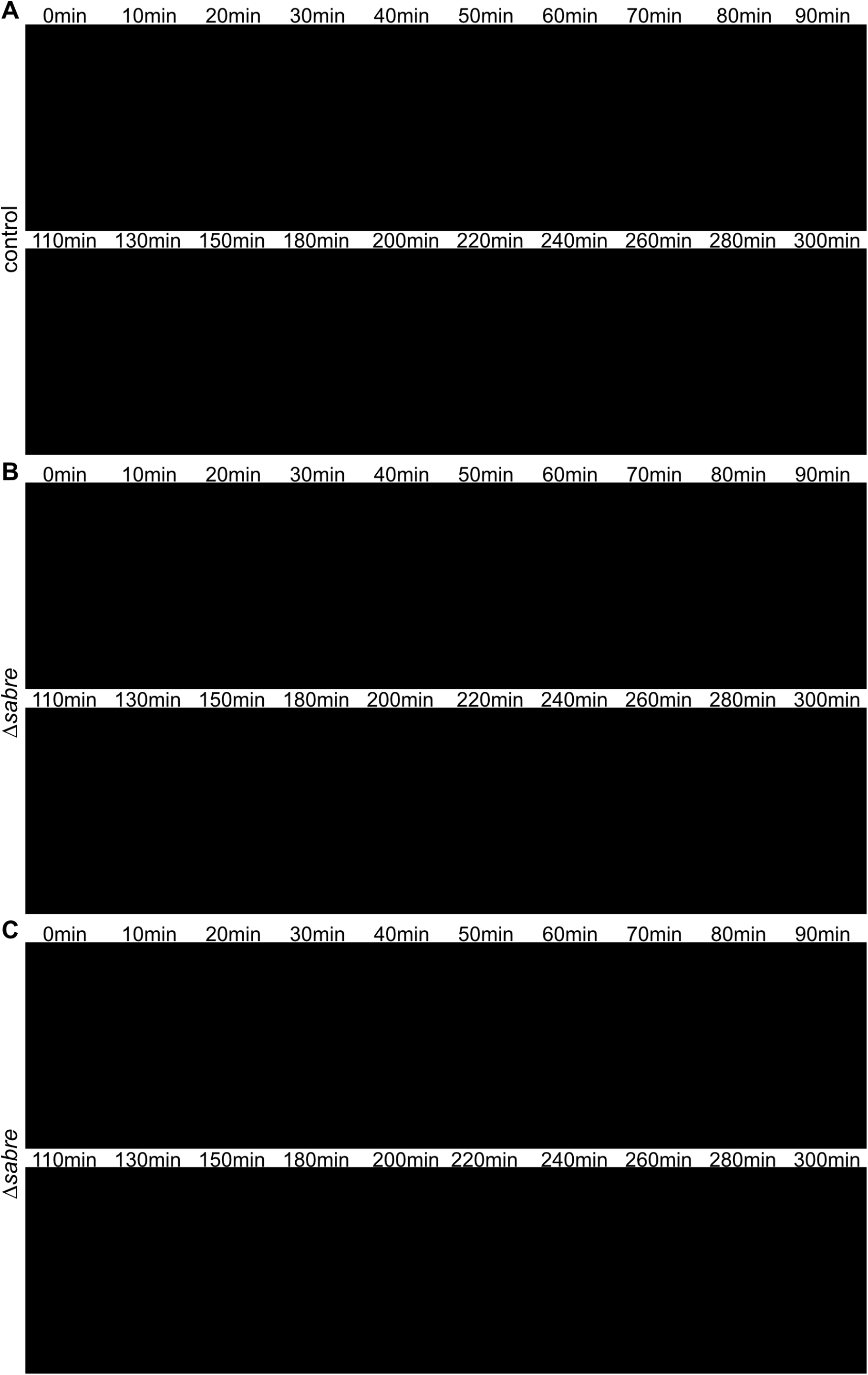
SABRE influences the ER, impacting nuclear movement and ER morphology at the nascent cell plate. (A) ER behavior in control cells during cell plate formation and nuclear migration after mitosis. Time points and arrows in blue indicate cell division and the initial apical nuclear movement (phase I), orange indicates basal nuclear movement (phase II), and black text accompanied with white arrows in the image indicate apical nuclear movement with polarized growth (phase III). (B-C) ER behavior in two example *Δsabre* cells. As in (A), the initial time points are within 10 minutes of nuclear envelope formation. Magenta arrowheads indicate ER buckling at the nascent cell plate. Red arrowheads indicate exaggerated basal nuclear movement. Green arrowheads indicate abnormally close contact between the nucleus and the nascent cell plate during the phase I apical movement. Scale bars, 5 μm. Also see Video 10.

**Figure 7 – figure supplement 1.**
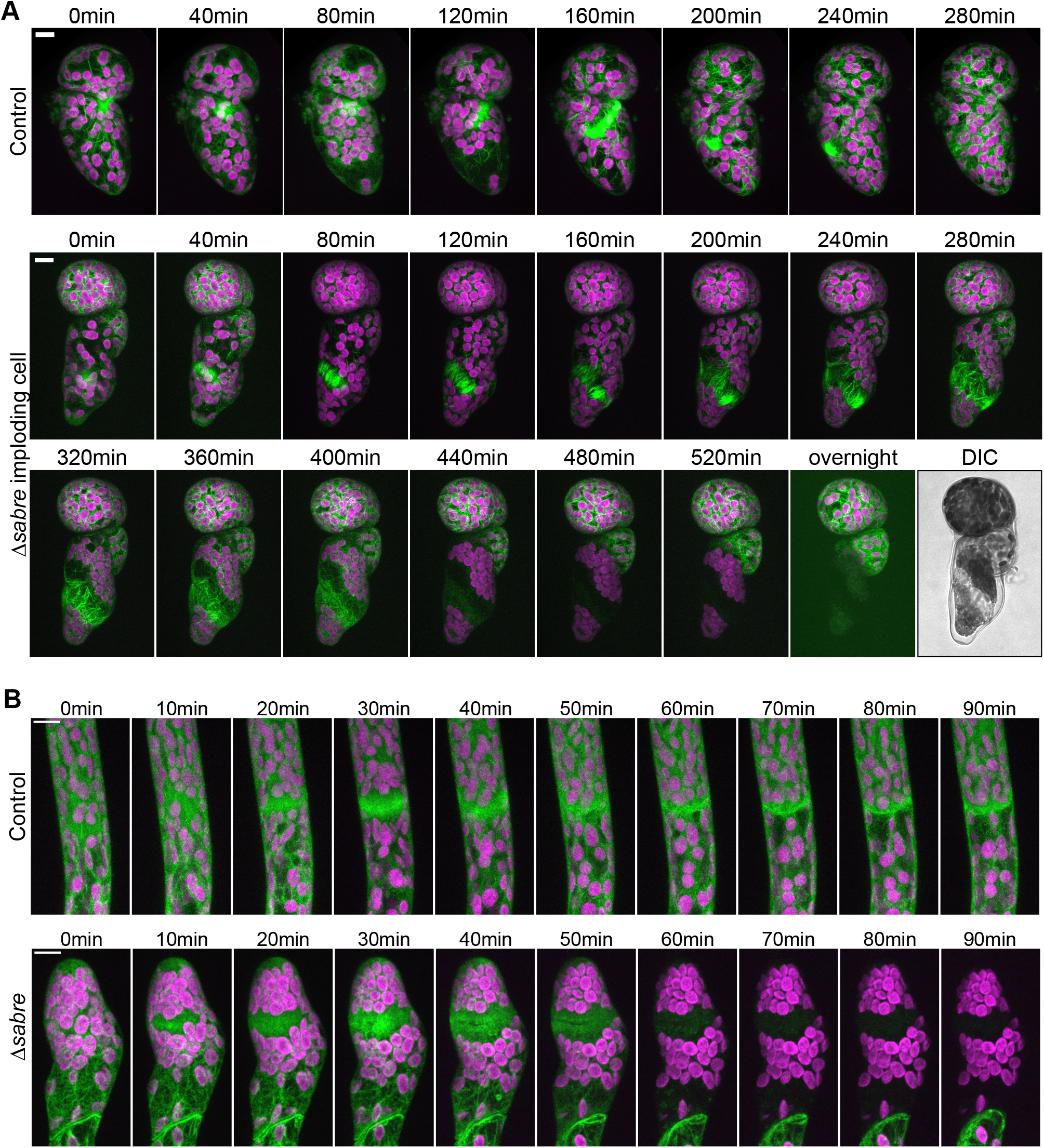
Phragmoplast microtubule and actin behavior during cytokinesis failures in *Δsabre* that resulted in cell lysis during cell division with no brown material deposition. (A) Time lapse imaging of GFP-tubulin (green) and chlorophyll autofluorescence (magenta) in 4-day old plants regenerated from protoplasts. Images are maximum projections of confocal Z-stacks. Top, a control cell dividing successfully. Bottom, a *Δsabre* cell exhibiting phragmoplast microtubule disorganization and rapid cell lysis. DIC image in B depicts the dead cell at the end of the movie. Scale bars, 10 μm. (B) Actin labeled with Lifeact-GFP (green) and chlorophyll autofluorescence (magenta) during cell division. Scale bars, 10 μm. Top, a control cell dividing successfully. Bottom, a *Δsabre* cell that lysed during cell plate formation.

